# A flexible pipeline combining clustering and correction tools for prokaryotic and eukaryotic metabarcoding

**DOI:** 10.1101/717355

**Authors:** Miriam I. Brandt, Blandine Trouche, Laure Quintric, Patrick Wincker, Julie Poulain, Sophie Arnaud-Haond

## Abstract

Environmental metabarcoding is an increasingly popular tool for studying biodiversity in marine and terrestrial biomes. With sequencing costs decreasing, multiple-marker metabarcoding, spanning several branches of the tree of life, is becoming more accessible. However, bioinformatic approaches need to adjust to the diversity of taxonomic compartments targeted as well as to each barcode gene specificities. We built and tested a pipeline based on Illumina read correction with DADA2 allowing analyzing metabarcoding data from prokaryotic (16S) and eukaryotic (18S, COI) life compartments. We implemented the option to cluster Amplicon Sequence Variants (ASVs) into Operational Taxonomic Units (OTUs) with swarm v2, a network-based clustering algorithm, and to further curate the ASVs/OTUs based on sequence similarity and co-occurrence rates using a recently developed algorithm, LULU. Finally, flexible taxonomic assignment was implemented via Ribosomal Database Project (RDP) Bayesian classifier and BLAST. We validate this pipeline with ribosomal and mitochondrial markers using eukaryotic mock communities and 42 deep-sea sediment samples. The results show that ASVs, reflecting genetic diversity, may not be appropriate for alpha diversity estimation of organisms fitting the biological species concept. The results underline the advantages of clustering and LULU-curation for producing more reliable metazoan biodiversity inventories, and show that LULU is an effective tool for filtering metazoan molecular clusters, although the minimum identity threshold applied to co-occurring OTUs has to be increased for 18S. The comparison of BLAST and the RDP Classifier underlined the potential of the latter to deliver very good assignments, but highlighted the need for a concerted effort to build comprehensive, ecosystem-specific, databases adapted to the studied communities.

## Introduction

High-throughput sequencing (HTS) technologies are revolutionizing the way we assess biodiversity. By producing millions of DNA sequences per sample, HTS allows broad taxonomic biodiversity surveys through metabarcoding of bulk DNA from complex communities or from environmental DNA (eDNA) directly extracted from soil, water, and air samples. First developed to unravel cryptic and uncultured prokaryotic diversity, metabarcoding methods have been extended to eukaryotes as powerful, non-invasive tools, allowing detection of a wide range of taxa in a rapid, cost-effective way using a variety of sample types [1]–[4]. In the last decade, these tools have been used to describe past and present biodiversity in terrestrial [5]–[9], freshwater [10]–[14], and marine [15]–[22] environments.

As every new technique brings on new challenges, a number of studies have put considerable effort into delineating critical aspects of metabarcoding protocols to ensure robust and reproducible results (see Fig.1 in [23]). Recent studies have addressed many issues regarding sampling methods [24], contamination risks [25], DNA extraction protocols [26]–[28], amplification biases and required PCR replication levels [29]–[31]. Similarly, computational pipelines, through which molecular data are transformed into ecological inventories of putative taxa, have also been in constant improvement. PCR-generated errors and sequencing errors are major bioinformatic challenges for metabarcoding pipelines, as they can strongly bias biodiversity estimates [32], [33]. A variety of tools have thus been developed for quality-filtering amplicon data to remove erroneous reads and improve the reliability of Illumina-sequenced metabarcoding inventories [33]–[35]. Studies that evaluated bioinformatic processing steps have generally found that sequence quality-filtering parameters and clustering thresholds most strongly affect molecular biodiversity inventories, resulting in considerable variation during data analysis[26], [36]–[38].

**Figure 1.**
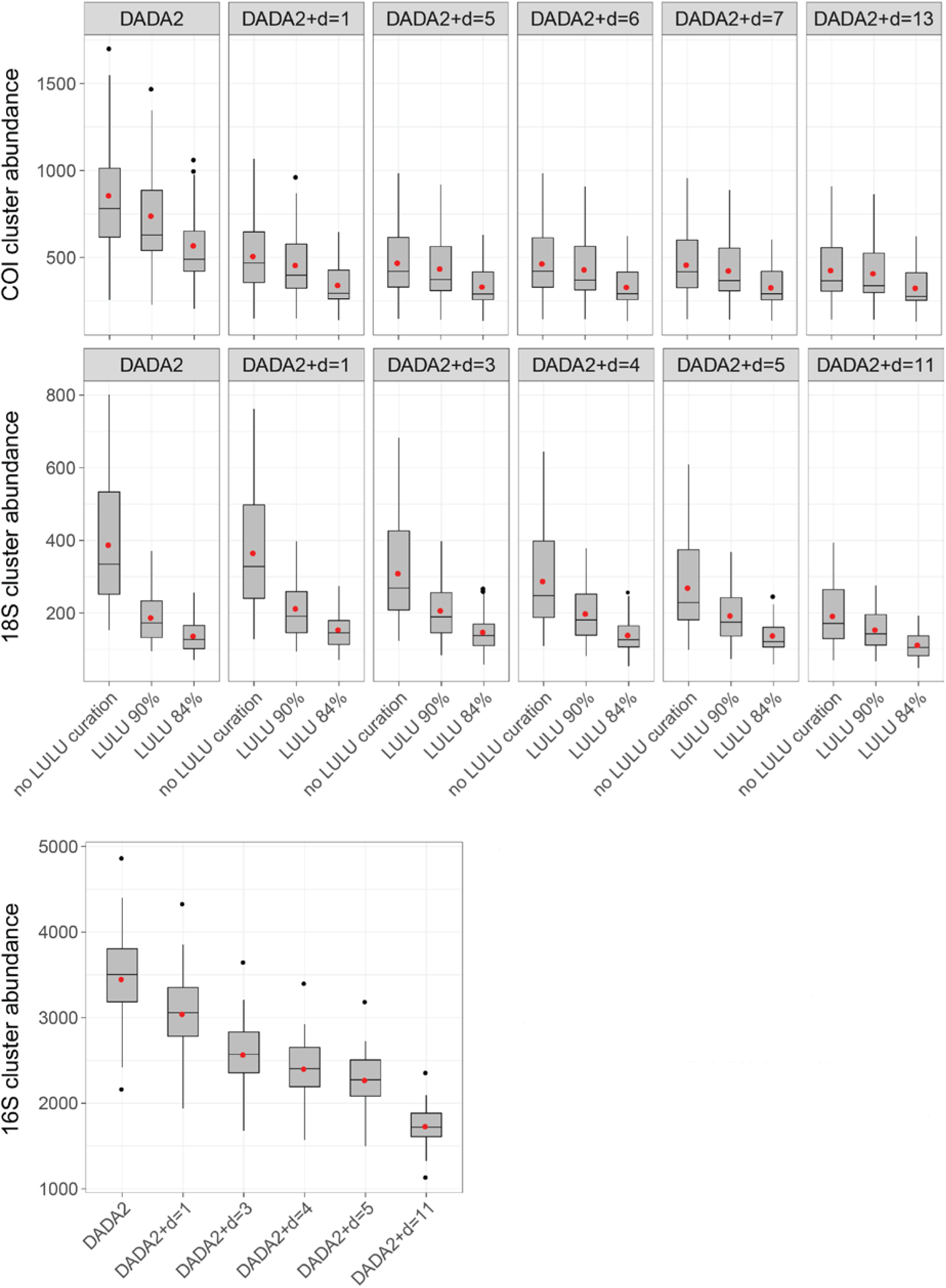
Number of COI, 18S, and 16S clusters detected in sediment of 14 deep-sea sites with the DADA2 metabarcoding pipeline, with and without swarm-clustering at different *d* values, and with and without LULU curation at 84% and 90% *minimum match*. Cluster abundance was obtained after rarefaction to minimal sequencing depth. Boxplots represent medians with first and third quartiles. Red dots indicate means.

There were historically two main reasons for clustering sequences into Operational Taxonomic Units (OTUs). The first was to limit the bias due to PCR and sequencing errors (and to some extent intra-individual variability linked to the existence of pseudogenes) by clustering erroneous sequences with error-free target sequences. The second was to delineate OTUs as clusters of homologous sequences (by grouping the alleles/haplotype at the same locus) that would best fit a “species level”, i.e. the Operational Taxonomic Units defined using a classical phenetic proxy [39]. Recent bioinformatic algorithms alleviate the influence of errors and intraspecific variability in metabarcoding datasets. First, amplicon-specific error correction methods, commonly used to correct sequences produced by pyrosequencing [32], have now become available for Illumina-sequenced data. Introduced in 2016, DADA2 effectively corrects Illumina sequencing errors and has quickly become a widely used tool, particularly in the microbial world, producing more accurate biodiversity inventories and resolving fine-scale genetic variation by defining Amplicon Sequence Variants (ASVs) [40], [41]. Second, LULU is a recently developed curation algorithm designed to filter out spurious clusters, originating from PCR and sequencing errors, or intra-individual variability (pseudogenes, heteroplasmy), based on their similarity and co-occurrence rate with more abundant clusters, allowing obtaining curated datasets while avoiding arbitrary abundance filters [42]. The authors validated their approach on metabarcoding of plants using ITS2 (nuclear ribosomal internal transcribed spacer region 2) and evaluated it on several pipelines. Their results show that ASV definition with DADA2, subsequent clustering to address intraspecific variation, and final curation with LULU is the safest pathway for producing reliable and accurate metabarcoding data. The authors concluded that their validation on plants is relevant to other organism groups and other markers, while recommending future validation of LULU on mock communities as LULU’s minimum match parameter may need to be adjusted to less variable marker genes.

The impact of errors being strongly decreased by correction algorithms such as DADA2 and LULU, the relevance of clustering sequences into OTUs is now being debated. Indeed, after presenting their new algorithm on prokaryotic communities, the authors of DADA2 proposed that the reproducibility and comparability of ASVs across studies challenge the need for clustering sequences, as OTUs have the disadvantage of being study-specific and defined using arbitrary thresholds [43]. However, clustering sequences may still be necessary in metazoan datasets, where very distinct levels of intraspecific polymorphism can exist in the same gene region among taxa due to both evolutionary and biological specificity [44], [45]. ASV-based inventories will thus be biased in favour of taxa with high levels of intraspecific diversity, even though the latter are not necessarily the most abundant ones [46]. Such bias in biodiversity inventories based on ASVs is likely to be magnified in presence-absence metabarcoding datasets, commonly used for metazoan communities [5]. Similarly, imposing a “universal” clustering threshold on metabarcoding datasets is also introducing bias, penalizing groups with lower interspecific divergence, and overestimating species diversity in groups with higher interspecific divergence. However, this can be alleviated with tools such as swarm v2, a single-linkage clustering algorithm [47]. Based on network theory, swarm v2 aggregates sequences iteratively and locally around seed sequences and determines coherent groups of sequences, independent of amplicon input order, allowing highly scalable and fine-scale clustering. Finally, it is widely recognized that homogeneous entities sharing a set of evolutionary and ecological properties, i.e. species [48], [49], sometimes referred to “ecotypes” for prokaryotes [50], [51], represent a fundamental category of biological organization that is the cornerstone of most ecological and evolutionary theories and empirical studies. Maintaining ASV information for feeding databases and cross-comparing studies is not incompatible with their clustering into OTUs, and this choice depends on the purpose of the study, i.e. providing a census of the extent and distribution of genetic polymorphism for a given gene, or a census of biodiversity to be used and manipulated in ecological or evolutionary studies.

Here we evaluate DADA2 and LULU, using them alone and in combination with swarm v2, to assess the performance of these new tools for metabarcoding of metazoan communities. Using both mitochondrial COI [52] and the V1-V2 region of 18S ribosomal RNA (rRNA) [16], we evaluated the need for clustering and the effectiveness of LULU curation to select pipeline parameters delivering the most accurate resolution of two deep-sea mock communities. We then test the different bioinformatic tools on a deep-sea sediment dataset in order to select an optimal trade-off between inflating biodiversity estimates and loosing rare biodiversity. As a baseline for comparison, and in the perspective of the joint study of metazoan and microbial taxa, we also analyzed the 16S V4-V5 rRNA barcode on these natural samples [53].

Our objectives were to (1) discuss the use of ASV vs OTU-centered datasets depending on taxonomic compartment and study objectives, and (2) determine the most adequate swarm-clustering and LULU curation thresholds that avoid inflating biodiversity estimates while retaining rare biodiversity.

## Methods

### 1 Preparation of samples

#### Mock communities

Genomic-DNA mass-balanced metazoan mock communities (5 ng/μL) were prepared using standardized 10 ng/μL DNA extracts of ten deep-sea specimens belonging to five taxonomic groups (Polychaeta, Crustacea, Anthozoa, Bivalvia, Gastropoda; Table S1). Specimen DNA was extracted using a CTAB extraction protocol, from muscle tissue or from whole polyps in the case of cnidarians. The mock communities differed in terms of ratios of total genomic DNA from each species, with increased dominance of three species and secondary species DNA input decreasing from 3% to 0.7%. We individually barcoded the species present in the mock communities: PCRs of both target genes were performed using the same primers as the ones used in metabarcoding (see below). The PCR reactions (25 μL final volume) contained 2 μL DNA template with 0.5 μM concentration of each primer, 1X Phusion Master Mix, and an additional 1 mM MgCl2 for COI. PCR amplifications (98 °C for 30 s; 40 cycles of 10 s at 98 °C, 45 s at 48 °C (COI) or 57 °C (18S), 30 s at 72 °C; and 72 °C for 5 min) were cleaned up with ExoSAP (Thermo Fisher Scientific, Waltham, MA, USA) and sent to Eurofins (Eurofins Scientific, Luxembourg) for Sanger sequencing. The barcode sequences obtained for all mock specimens were added to the databases used for taxonomic assignments of metabarcoding datasets, and were submitted on Genbank under accession numbers MN826120-MN826130 and MN844176-MN844185.

#### Environmental DNA

Sediment cores were collected from fourteen deep-sea sites ranging from the Arctic to the Mediterranean during various cruises (Table S2). Sampling was carried out with a multicorer or with a remotely operated vehicle. Three tube cores were taken at each sampling station (GPS coordinates in Table S2). The latter were sliced into depth layers that were transferred into zip-lock bags, homogenised, and frozen at −80°C on board before being shipped on dry ice to the laboratory. The first layer (0-1 cm) was used in the present study. DNA extractions were performed using approximately 10 g of sediment with the PowerMax Soil DNA Isolation Kit (Qiagen, Hilden, Germany). To increase the DNA yield, the elution buffer was left on the spin filter membrane for 10 min at room temperature before centrifugation. The ~5 mL extract was then split into three parts, one of which was kept in screw-cap tubes for archiving purposes and stored at −80°C. For the four field controls, the first solution of the kit was poured into the control zip-lock bag, before following the usual extraction steps. For the two negative extraction controls, a blank extraction (adding nothing to the bead tube) was performed alongside sample extractions.

### 2 Amplicon library construction and high-throughput sequencing

Two primer pairs were used to amplify the mitochondrial COI and the 18S V1-V2 rRNA barcode genes specifically targeting metazoans, and one pair of primer was used to amplify the prokaryote 16S V4-V5 region. PCR amplifications, library preparation, and sequencing were carried out at Genoscope (Evry, France) as part of the eDNAbyss project.

#### Eukaryotic 18S V1-V2 rRNA gene amplicon generation

Amplifications were performed with the Phusion High Fidelity PCR Master Mix with GC buffer (Thermo Fisher Scientific, Waltham, MA, USA) and the SSUF04 (5’- GCTTGTCTCAAAGATTAAGCC-3’) and SSUR22mod (5’- CCTGCTGCCTTCCTTRGA-3’) primers [16], preferentially targeting metazoans, the primary focus of this study. The PCR reactions (25 μL final volume) contained 2.5 ng or less of DNA template with 0.4 μM concentration of each primer, 3% of DMSO, and 1X Phusion Master Mix. PCR amplifications (98 °C for 30 s; 25 cycles of 10 s at 98 °C, 30 s at 45 °C, 30 s at 72 °C; and 72 °C for 10 min) of all samples were carried out in triplicate in order to smooth the intra-sample variance while obtaining sufficient amounts of amplicons for Illumina sequencing.

#### Eukaryotic COI gene amplicon generation

Metazoan COI barcodes were generated using the mlCOIintF (5’- GGWACWGGWTGAACWGTWTAYCCYCC-3’) and jgHCO2198 (5’-TAIACYTCIGGRTGICCRAARAAYCA-3’) primers [52].Triplicate PCR reactions (20 μl final volume) contained 2.5 ng or less of total DNA template with 0.5 μM final concentration of each primer, 3% of DMSO, 0.175 mM final concentration of dNTPs, and 1X Advantage 2 Polymerase Mix (Takara Bio, Kusatsu, Japan). Cycling conditions included a 10 min denaturation step followed by 16 cycles of 95 °C for 10 s, 30s at 62°C (−1°C per cycle), 68 °C for 60 s, followed by 15 cycles of 95 °C for 10 s, 30s at 46°C, 68 °C for 60 s and a final extension of 68 °C for 7 min.

#### Prokaryotic 16S rRNA gene amplicon generation

Prokaryotic barcodes were generated using 515F-Y (5’- GTGYCAGCMGCCGCGGTAA-3’) and 926R (5’- CCGYCAATTYMTTTRAGTTT-3’) 16S-V4V5 primers [53]. Triplicate PCR mixtures were prepared as described above for 18S-V1V2, but cycling conditions included a 30 s denaturation step followed by 25 cycles of 98 °C for 10 s, 53 °C for 30 s, 72 °C for 30 s, and a final extension of 72 °C for 10 min.

#### Amplicon library preparation

PCR triplicates were pooled and PCR products purified using 1X AMPure XP beads (Beckman Coulter, Brea, CA, USA) clean up. Aliquots of purified amplicons were run on an Agilent Bioanalyzer using the DNA High Sensitivity LabChip kit (Agilent Technologies, Santa Clara, CA, USA) to check their lengths and quantified with a Qubit fluorimeter (Invitrogen, Carlsbad, CA, USA). One hundred nanograms of pooled amplicon triplicates were directly end-repaired, A-tailed and ligated to Illumina adapters on a Biomek FX Laboratory Automation Workstation (Beckman Coulter, Brea, CA, USA). Library amplification was performed using a Kapa Hifi HotStart NGS library Amplification kit (Kapa Biosystems, Wilmington, MA, USA) with the same cycling conditions applied for all metagenomic libraries and purified using 1X AMPure XP beads.

#### Sequencing library quality control

Amplicon libraries were quantified by Quant-iT dsDNA HS assay kits using a Fluoroskan Ascent microplate fluorometer (Thermo Fisher Scientific, Waltham, MA, USA) and then by qPCR with the KAPA Library Quantification Kit for Illumina Libraries (Kapa Biosystems, Wilmington, MA, USA) on an MxPro instrument (Agilent Technologies, Santa Clara, CA, USA). Library profiles were assessed using a high-throughput microfluidic capillary electrophoresis system (LabChip GX, Perkin Elmer, Waltham, MA, USA).

#### Sequencing procedures

Library concentrations were normalized to 10 nM by addition of 10 mM Tris-Cl (pH 8.5) and applied to cluster generation according to the Illumina Cbot User Guide (Part # 15006165). Amplicon libraries are characterized by low diversity sequences at the beginning of the reads due to the presence of the primer sequence. Low-diversity libraries can interfere in correct cluster identification, resulting in a drastic loss of data output. Therefore, loading concentrations of libraries were decreased (8–9 pM instead of 12–14 pM for standard libraries) and PhiX DNA spike-in was increased (20% instead of 1%) in order to minimize the impacts on the run quality.

Libraries were sequenced on HiSeq2500 (System User Guide Part # 15035786) instruments (Illumina, San Diego, CA, USA) in a 250 bp paired-end mode.

### 3 Bioinformatic analyses

All bioinformatic analyses were performed using a Unix shell script on a home-based cluster (DATARMOR, Ifremer), available on Gitlab (https://gitlab.ifremer.fr/abyss-project/). The mock communities were analysed alongside the natural samples, and used to validate the metabarcoding pipeline in terms of detection of correct species and presence of false-positives. The details of the pipeline, along with specific parameters used for all three metabarcoding markers are listed in Table S3.

#### Reads preprocessing

Our multiplexing strategy relies on ligation of adapters to amplicon pools, meaning that contrary to libraries produced by double PCR, the reads in each paired sequencing run can be forward or reverse. DADA2 correction is based on error distribution differing between R1 and R2 reads. We thus developed a custom script (abyss-preprocessing in abyss-pipeline) allowing separating forward and reverse reads in each paired run and reformatting the outputs to be compatible with DADA2. Briefly, the script uses cutadapt v1.18 to detect and remove primers, while separating forward and reverse reads in each paired sequence file to produce two pairs of sequence files per sample named R1F/R2R and R2F/R1R. Cutadapt parameters (Table S3) were set to require an overlap over the full length of the primer (default: 3 nt), with 2-4 nt mismatches allowed for ribosomal loci, and 7 nt mismatches allowed for COI (default: 10%). Each identified forward and reverse read is then renamed which the correct extension (/1 and /2 respectively), which is a requirement for DADA2 to recognize the pairs of reads. Each pair of renamed sequence files is then re-paired with BBMAP Repair v38.22 in order to remove singleton reads (non-paired reads). Optionally, sequence file names can also be renamed if necessary using a CSV correspondence file.

#### Read correction, amplicon cluster generation and taxonomic assignment

Pairs of Illumina reads were corrected with DADA2 v.1.10 [40] following the online tutorial for paired-end HiSeq data (https://benjjneb.github.io/dada2/bigdata_paired.html). Reads were filtered and trimmed with the filterAndTrim function and all reads containing ambiguous bases removed. The parameters were set based on tutorial recommendations and trimming lengths were adjusted based on sequence quality profiles, so that Q-scores remained above 30 (truncLen at 220 for 18S and 16S, 200 for COI, maxEE at 2, truncQ at 11, maxN at 0).

The error model was calculated for forward and reverse reads (R1F/R2R pairs and then R2F/R1R pairs) with learnErrors based on 100 million randomly chosen bases (default), and reads were dereplicated using derepFastq. After read correction with the dada function, forward and reverse reads were merged with a minimum overlap of 12 nucleotides, allowing no mismatches (default). The amplicons were then filtered by size. The size range was set to 330-390 bp for the 18S SSU rRNA marker gene, 300-326 bp for the COI marker gene, and 350-390 bp for the 16S rRNA marker gene.

Chimeras were removed with removeBimeraDenovo and ASVs were taxonomically assigned via the RDP naïve Bayesian classifier method, the default assignment method implemented in DADA2. A second taxonomic assignment method was optionally implemented in the pipeline, allowing assigning ASVs using BLAST+ (Basic Local Alignment Search Tool v2.6.0) based on minimum similarity and minimum coverage (-perc_identity 70 and ‒qcov_hsp 80). An initial test implementing BLASTn+ to assign taxonomy only to the COI dataset using a 96% percent identity threshold led to the exclusion of the majority of the clusters. Given observed inter-specific mitochondrial DNA divergence levels of up to 30% within a same polychaete genus [54] or among some closely related deep-sea shrimp species [55], and considering our interest in the identities of multiple, largely unknown taxa in poorly characterized communities, more stringent BLAST thresholds were not implemented at this stage. The Silva132 reference database was used for the 16S and 18S SSU rRNA marker genes [56], and MIDORI-UNIQUE [57] was used for COI. The databases were downloaded from the DADA2 website (https://benjjneb.github.io/dada2/training.html) and from the FROGS website (http://genoweb.toulouse.inra.fr/frogs_databanks/assignation/). Finally, to evaluate the effect of clustering, ASV tables produced by DADA2 were clustered with swarm v2 [47] at d=1,3,4,5 and 11 for 18S and 16S, and d=1,5,6,7, and 13 for COI in FROGS (http://frogs.toulouse.inra.fr/) [58]. Resulting OTUs were taxonomically assigned via RDP and BLAST+ using the databases stated above.

Molecular clusters were refined in R v.3.5.1 [59]. A blank correction was made using the decontam package v.1.2.1 [60], removing all clusters that were prevalent (more frequent) in negative control samples. ASV/OTU tables were refined taxonomically based on their RDP or BLAST taxonomy. For both assignment methods, unassigned clusters were removed. Non-target 18S and COI clusters (bacterial, non-metazoan) as well as all clusters with a terrestrial assignment (taxonomic groups known to be terrestrial-only, such as Insecta, Arachnida, Diplopoda, Amphibia, terrestrial mammals, Stylommatophora, Aves, Onychophora, Succineidae, Cyclophoridae, Diplommatinidae, Megalomastomatidae, Pupinidae, Veronicellidae) were removed. Samples were checked to ensure that a minimum of 10,000 metazoan reads were left after refining. Finally, as tag-switching is always to be expected in multiplexed metabarcoding analyses [61], an abundance renormalization was performed to remove spurious positive results due to reads assigned to the wrong sample [62], the original R script being available at https://github.com/metabarpark/R_scripts_metabarpark.

To test LULU curation [42], refined 18S and COI ASVs/OTUs were curated with LULU v.0.1 following the online tutorial (https://github.com/tobiasgf/lulu). The LULU algorithm detects erroneous clusters by comparing their sequence similarities and co-occurrence rate with more abundant (“parent”) clusters. LULU was tested with a minimum relative co-occurrence of 0.90, using a minimum similarity threshold (minimum match) at 84% (default) and slightly higher at 90%, following recommendations of the authors for less variable loci than ITS.

The vast majority of prokaryotes usually show low levels (< 1% divergence) of intra genomic variability for the 16S SSU rRNA gene [63], [64]. These low intragenomic divergence levels can be efficiently removed with swarm clustering at d=1. Although LULU curation may still be useful to merge redundant phylotypes in specific cases such as haplotype network analyses, this was not tested in this study. Indeed, parallelization not being currently available for LULU curation, the richness of prokaryote communities implied an unrealistic calculation time, even on a powerful cluster (e.g. LULU curation was at 20-40% after 4 days of calculation on our cluster).

### 4 Statistical analyses

Sequence tables were analysed using R with the packages phyloseq v1.22.3 [65] following guidelines on online tutorials (http://joey711.github.io/phyloseq/tutorials-index.html), and vegan v2.5.2 [66]. The datasets were normalized by rarefaction to their common minimum sequencing depth, before analysis of mock communities and natural samples.

To evaluate the functionality of the pipeline with the mock communities, taxonomically assigned metazoan clusters were considered as derived from one of the ten species used for the mock communities when the assignment delivered the corresponding species, genus, family, or class. Clusters not fitting the expected taxa were labelled as ‘Others’. Apart from PCR errors, these non-target clusters may also originate from contamination by external DNA from associated microfauna, or gut content in the case of whole polyps used for cnidarians.

Alpha diversity detected using each pipeline in the natural samples was evaluated with the number of observed target-taxa in the rarefied datasets via analyses of variance (ANOVA) on generalized linear models based on quasipoisson distribution models. Homogeneity of multivariate dispersions were verified with the betapart package v.1.5.1 [67]. Beta-diversity patterns were visualised via Principal Coordinates Analyses (PCoA), using Jaccard dissimilarities for metazoans and Bray-Curtis dissimilarities for prokaryotes. The effect of site and LULU curation on community composition was tested by means of PERMANOVA, using the function adonis2 (vegan), with the same dissimilarities as in PCoAs, and permuting 999 times. Finally, BLAST and RDP taxonomic assignments of the mock samples and the global dataset were compared at the most adequate pipeline settings for each locus. BLAST-refined (minimum identity at 70%) and RDP-refined (minimum phylum bootstrap at 80%) datasets were compared on ASV-level for prokaryotes, and OTU-level for metazoans (swarm d=3, LULU at 84% for COI and 90% for 18S). As trials on MIDORI-UNIQUE resulted in very poor performance of RDP for COI (assignments belonging mostly to Insecta), the comparison was performed with MIDORI-UNIQUE subsampled to marine taxa only.

## Results

### 1 Alpha diversity in mock communities

A number of 2 million (18S) and 1.5 million (COI) raw reads were obtained from the two mock communities (Table S4).After refining, these numbers were decreased to 1.3 million for 18S and 0.7 million for COI.

Seven out of ten mock species were recovered in the 18S dataset and all species were detected in the COI dataset (Table 1), even with minimum relative DNA abundance levels as low as 0.7% (Mock 5). Taxonomically unresolved species were correctly assigned up to their common family or class level. Dominant species generally produced more reads in both the clustered and non-clustered datasets (Table S6). When ASVs were clustered with swarm v2, this generally led to a slight loss of taxonomic resolution: Chorocaris sp. was not detected in Mock 5 for 18S at d > 1, and the two bivalves P. kilmeri and C. regab were taxonomically misidentified for COI at d ≥ 1.

**Table 1.** Number of ASVs/OTUs detected per species in the mock communities using different bioinformatic pipelines. White cells indicate an exact match with the number of OTUs expected, grey cells indicate a number of OTUs differing by ±3 from the number expected, and dark grey cells indicate a number of OTUs >3 from the one expected.

| 18S | DADA2 | DADA2<br>+LULU<br>90% | DADA2+<br>LULU<br>84% |  | DADA2+swarm<br>d1/d3/d4/d5/d11 | DADA2+swarm<br>d1/d3/d4/d5/d11 +<br>LULU 90% | DADA2+swarm<br>d1/d3/d4/d5/d11 +<br>LULU 84% |
| --- | --- | --- | --- | --- | --- | --- | --- |
| **Mock 3** |  |  |  |  |  |  |  |
| Alcyonacea;*A.arbuscula* | 64 | 1 | 1 | Alcyonacea;*A.arbuscula* | 29/11/9/7/6 | 1/1/1/1/1 | 1/1/1/1/1 |
| Caryophylliidae;*D.dianthus* | 2 | 1 | 1 | Caryophylliidae;*D.dianthus* | 2/2/1/1/1 | 1/1/1/1/1 | 1/1/1/1/1 |
| *Alvinocaris muricola* | 2 | 1 | 1 | *Alvinocaris muricola* | 2/1/1/1/1 | 1/1/1/1/1 | 1/1/1/1/1 |
| *Chorocaris* sp. | 1 | 0 | 0 | *Chorocaris* sp. | 2/1/1/1/1 | 0/0/0/0/0 | 0/0/0/0/0 |
| *Munidopsis* sp. | 6 | 1 | 1 | *Munidopsis* sp. | 5/4/3/3/2 | 1/1/1/1/1 | 1/1/1/1/1 |
| Gastropoda;*Paralepetopsis* sp. | 1 | 1 | 0 | Gastropoda;*Paralepetopsis* sp. | 1/1/1/1/1 | 1/1/1/1/1 | 0/0/0/0/0 |
| Vesicomidae;*P. kilmeri*/*C. regab*/*V. gigas* | 8 | 1 | 1 | Bivalvia;*P. kilmeri*/*C. regab*/*V. gigas* | 5/4/4/4/2 | 1/2/2/2/1 | 1/1/1/1/1 |
| Polychaeta;*E.norvegica* | 8 | 3 | 2 | Polychaeta;*E.norvegica* | 5/4/4/4/3 | 3/2/2/2/2 | 2/1/2/2/2 |
| Others | 3 | 3 | 2 | Others | 4/4/4/4/4 | 2/2/2/2/3 | 2/2/2/2/2 |
| **Mock 5** |  |  |  |  |  |  |  |
| Alcyonacea;*A.arbuscula* | 54 | 1 | 1 | Alcyonacea;*A.arbuscula* | 28/11/9/7/6 | 1/1/1/1/1 | 1/1/1/1/1 |
| Caryophylliidae;*D.dianthus* | 1 | 1 | 1 | Caryophylliidae;*D.dianthus* | 1/1/1/1/1 | 1/1/1/1/1 | 1/1/1/1/1 |
| *Alvinocaris muricola* | 1 | 1 | 1 | *Alvinocaris muricola* | 1/1/1/1/1 | 1/1/1/1/1 | 1/1/1/1/1 |
| *Chorocaris* sp. | 1 | 0 | 0 | *Chorocaris* sp. | 1/0/0/0/0 | 0/0/0/0/0 | 0/0/0/0/0 |
| *Munidopsis* sp. | 4 | 1 | 1 | *Munidopsis* sp. | 4/3/3/3/2 | 1/1/1/1/1 | 1/1/1/1/1 |
| Gastropoda;*Paralepetopsis* sp. | 1 | 1 | 0 | Gastropoda;*Paralepetopsis* sp. | 1/1/1/1/1 | 1/1/1/1/1 | 0/0/0/0/0 |
| Vesicomidae;*P. kilmeri*/*C. regab*/*V. gigas* | 5 | 1 | 1 | Bivalvia;*P. kilmeri*/*C. regab*/*V. gigas* | 5/3/3/3/2 | 1/1/1/1/1 | 1/1/1/1/1 |
| Polychaeta;*E.norvegica* | 11 | 3 | 2 | Polychaeta;*E.norvegica* | 5/4/4/4/3 | 3/2/2/2/1 | 2/1/2/2/2 |
| Others | 4 | 3 | 2 | Others | 3/4/4/4/2 | 4/2/2/2/1 | 4/2/2/2/3 |
| COI | DADA2 | DADA2<br>+LULU<br>90% | DADA2+<br>LULU<br>84% |  | DADA2+swarm<br>d1/d5/d6/d7/d13 | DADA2+swarm<br>d1/d5/d6/d7/d13 +<br>LULU 90% | DADA2+swarm<br>d1/d5/d6/d7/d13 +<br>LULU 84% |
| **Mock 3** |  |  |  |  |  |  |  |
| *Acanella arbuscula* | 1 | 1 | 1 | *Acanella arbuscula* | 1/1/1/1/1 | 1/1/1/1/1 | 1/1/1/1/1 |
| Hexacorallia;*D.dianthus* | 3 | 3 | 3 | Hexacorallia;*D.dianthus* | 3/4/4/4/3 | 3/3/3/3/3 | 3/3/3/3/3 |
| *Alvinocaris* A. *muricola* | 26 | 2 | 2 | *Alvinocaris* A. *muricola* | 21/12/10/10/5 | 1/1/1/1/1 | 1/1/1/1/1 |
| *Chorocaris* sp. | 2 | 1 | 1 | *Chorocaris* sp. | 3/3/3/3/3 | 1/1/1/1/1 | 1/1/1/1/1 |
| *Munidopsis* sp. | 2 | 1 | 1 | *Munidopsis* sp. | 3/2/1/1/1 | 2/1/1/1/1 | 1/1/1/1/1 |
| Gastropoda;*Paralepetopsis* sp. | 8 | 2 | 3 | Gastropoda;*Paralepetopsis* sp. | 3/3/3/3/2 | 2/2/2/2/2 | 2/2/2/2/2 |
| *Phreagena kilmeri* | 2 | 1 | 1 | Bivalvia;*P. kilmeri* | 2/3/3/3/3 | 2/2/2/2/2 | 2/2/2/2/2 |
| Bivalvia;*C. regab* | 2 | 1 | 1 | Bivalvia;*C. regab* |  |  |  |
| *Vesicomya gigas* | 1 | 1 | 1 | *Vesicomya gigas* | 1/1/1/1/1 | 1/1/1/1/1 | 1/1/1/1/1 |
| Polychaeta;*E.norvegica* | 3 | 2 | 1 | *Eunice norvegica* | 2/1/1/1/1 | 2/1/1/1/1 | 1/1/1/1/1 |
| Others | 7 | 6 | 6 | Others | 3/3/3/3/4 | 4/5/5/5/5 | 5/5/5/5/5 |
| **Mock 5** |  |  |  |  |  |  |  |
| *Acanella arbuscula* | 1 | 1 | 1 | *Acanella arbuscula* | 1/1/1/1/1 | 1/1/1/1/1 | 1/1/1/1/1 |
| Hexacorallia;*D.dianthus* | 3 | 3 | 3 | Hexacorallia;*D.dianthus* | 3/3/3/3/3 | 3/3/3/3/3 | 3/3/3/3/3 |
| *Alvinocaris* A. *muricola* | 26 | 2 | 2 | *Alvinocaris* A. *muricola* | 21/12/10/10/5 | 1/1/1/1/1 | 1/1/1/1/1 |
| *Chorocaris* sp. | 1 | 1 | 1 | *Chorocaris* sp. | 2/2/2/2/2 | 1/1/1/1/1 | 1/1/1/1/1 |
| *Munidopsis* sp. | 2 | 1 | 1 | *Munidopsis* sp. | 2/2/1/1/1 | 1/1/1/1/1 | 1/1/1/1/1 |
| Gastropoda;*Paralepetopsis* sp. | 5 | 2 | 2 | Gastropoda;*Paralepetopsis* sp. | 3/2/2/2/2 | 2/2/2/2/2 | 2/2/2/2/2 |
| *Phreagena kilmeri* | 1 | 1 | 1 | Bivalvia;*P. kilmeri* | 2/2/2/2/2 | 2/2/2/2/2 | 2/2/2/2/2 |
| Bivalvia;*C. regab* | 2 | 1 | 1 | Bivalvia;*C. regab* |  |  |  |
| *Vesicomya gigas* | 1 | 1 | 1 | *Vesicomya gigas* | 1/1/1/1/1 | 1/1/1/1/1 | 1/1/1/1/1 |
| Polychaeta;*E.norvegica* | 3 | 2 | 1 | *Eunice norvegica* | 2/2/2/2/2 | 1/1/1/1/1 | 1/1/1/1/1 |
| Others | 6 | 5 | 4 | Others | 2/2/2/2/2 | 1/2/2/2/2 | 1/1/1/1/1 |

Clustering sequences with swarm v2 reduced the number of clusters produced per species, but some species still produced multiple OTUs even at d values as high as d=11 for 18S (A. arbuscula, Munidopsis sp., and E. norvegica) and d=13 for COI D. dianthus, A. muricola, Chorocaris sp., and Paralepetopsis sp.). Curating with LULU allowed reducing the number of clusters produced per species to nearly one for both loci, but the best results were obtained in datasets clustered at d > 1 for 18S and d ≥ 1 for COI. Moreover, LULU curation tended to decrease the number of non-target clusters (“Others”) (Table 1). In the clustered COI dataset, curating with LULU at 84% minimum match resulted in the most accurate detection of community composition, and this for all d values tested. However, curating with LULU the 18S data (ASVs or OTUs) led to the loss of one shrimp species (Chorocaris sp) when the minimum match parameter was at 90% and an additional species was lost (the limpet Paralepetopsis sp.) when this parameter was at 84%. LULU consistently merged the shrimp species Chorocaris sp with another shrimp species as the latter were always co-occurring in our mock samples.

### 2 Alpha-diversity patterns in natural samples

#### High-throughput sequencing results

A number of 44 million (18S), 33 million (COI) and 16 million (16S) reads were obtained from 42 sediment samples, 4 field controls, 2 extraction blanks, and 4-10 PCR blanks (Table S4). Two sediment samples failed amplification for the COI marker gene (PCT_FA_CT2_0_1 and CHR_CT1_0_1). For metazoans, less reads were retained after bioinformatic processing in negative controls (36% for 18S, 47% for COI) compared to true samples (~60% for 18S, ~70% for COI), while the opposite was observed for 16S (74% of reads retained in control samples against 53% in true samples). Negative control samples (field, extraction, and PCR controls) contained 2,186,230 (~8%) 18S reads, 1,015,700 (~4%) COI reads, and 2,618,729 (28%) 16S reads. These reads were mostly originating from the field controls for metazoans (48% for 18S, 55% for COI) and extractions controls for 16S (50%).

After blank correction, data refining, and abundance renormalization, rarefaction curves showed that a plateau was achieved for all samples in both clustered and non-clustered datasets, suggesting an overall sequencing depth adequate to capture the diversity present (Fig. S1). The final 18S datasets (with and without clustering at selected d values) contained 8.9-9.6 million marine metazoan reads in 42 sediment samples (Table S4), and comprised 57,661 ASVs and 19,504-44,948 OTUs (Table S6). The final COI datasets contained 4.5-6.9 million marine metazoan reads in 40 sediment samples, and comprised 78,785 ASVs and 44,684-64,669 OTUs. The 16S datasets contained from 6.6 to 6.7 million prokaryotic reads in 42 sediment samples, producing 56,577 ASVs and 41,746-14,631 OTUs.

#### Number of clusters among pipelines

The number of metazoan clusters detected in the deep-sea sediment samples varied significantly between bioinformatic pipelines chosen (, and also varied significantly among sites (Table 2). However, the pipeline effect was consistent across sites although mean cluster numbers detected per sample spanned a wide range in all loci (100-800 for 18S, 150-1,500 for COI datasets, and 1,500-5,000 for 16S, Fig. 1).

**Table 2.** Effect of pipeline and site on the number of metazoan and prokaryote clusters. Results of the analysis of variance (ANOVA) of the rarefied cluster richness for the three genes studied. Pairwise comparisons were performed with Tukey’s HSD tests. DS: Dada2+swarm; DSL: Dada2+swarm+LULU; d: swarm *d-value*. Significance codes: ***: p<0.001; **: p<0.01; *: p<0.05.

| LOCUS | F-value | p-value | Significant pairwise comparisons |
| --- | --- | --- | --- |
| **COI** |  |  |  |
| Pipeline | 123.13 | $p < 0.001$ | Dada2 > DS***; DS(d1) > DS(d13)***; |
| Site | 356.37 | $p < 0.001$ | Dada2 > DL***; DS > DSL 84%***; D(S)L 90% > D(S)L 84%*** |
| Pipeline x Site | 0.16 | $p > 0.05$ | DL > DSL***; DL 90% > DS*** |
| **18S V1-V2** |  |  |  |
| Pipeline | 129.16 | $p < 0.001$ | Dada2 > DS(d>1)***; DS(d1) > DS(d>1)***; DS(d11) < DS(d1-5)***; |
| Site | 154.52 | $p < 0.001$ | Dada2 > DL***; DS > DSL 84%***; D(S)L 90% > D(S)L 84%***; |
| Pipeline x Site | 0.49 | $p > 0.05$ | DL 84% < DS*** |
| **16S V4-V5** |  |  |  |
| Pipeline | 179.19 | $p < 0.001$ | Dada2 > DS***; |
| Site | 18.46 | $p < 0.001$ | DS(d1) > DS(d>1)***; DS(d11) < DS(d1-5)*** |
| Pipeline x Site | 0.06 | $p > 0.05$ | |

Expectedly, clustering significantly reduced the number of detected clusters per sample for all loci. Consistent to results observed in mock communities, clustering at d=1-13 resulted in comparable OTU numbers for COI, while significantly higher OTU numbers were obtained at d=1 than with d >1 for ribosomal loci (Fig. 1, Table 2). DADA2 detected on average 863 (SE=61) metazoan COI ASVs per sample, and clustering reduced this number to around 500, regardless the d-value. For ribosomal loci, clustering at d=3-5 reduced OTU numbers of around 25-30% compared to without clustering, while at d=11, cluster numbers were halved.

LULU curation of metazoan ASVs significantly decreased the number of clusters detected at both tested minimum match values (Table 2). For OTU datasets, the decrease was significant only when the minimum match parameter was at 84%. The effect of LULU curation was stronger at a lower minimum match value for both loci, as LULU curation at 90% of ASVs or OTUs resulted in significantly more clusters than when the minimum match was at 84% (Table 2). The effect of LULU curation of was also more pronounced for the 18S locus: LULU decreased by 31-65% the number of 18S ASVs/OTUs, compared to 7-33% for COI. LULU curation of ASVs or OTUs resulted in comparable cluster numbers in the 18S datasets, regardless the d-value used for clustering. For example, at 84% minimum match, LULU curation produced on average 137 ± 7 and 140 ± 8 clusters per sample after application on ASVs and OTUs (d=4) respectively. At 90%, these numbers were at 189 ± 11 and 200 ± 12 (Fig. 1). This was not the case for COI, where LULU curation of ASVs resulted in significantly more clusters (574 ± 38 at 84% and 742 ± 53 at 90%) than LULU curation of OTUs (334 ± 21 and 433 ± 31 for d=6). Looking at mean ASV and OTU numbers detected per phylum with each pipeline showed consistent effects of swarm clustering and LULU curation, but highlighted strong differences in the amount of intragenomic variation between taxonomic groups. For all loci investigated, some taxa displayed high ASV to OTU ratios, while others were hardly affected by clustering or LULU curation in terms of numbers of clusters detected (Fig S2).

### 3 Patterns of beta-diversity between pipelines

Community differences were visualized using PCoA ordinations (Jaccard and Bray-Curtis dissimilarities for metazoans and prokaryotes respectively) in clustered and non-clustered datasets (Fig. 2, Fig. S3). Expectedly, PERMANOVAs confirmed that sites differed significantly in terms of community structure, accounting from 45% to 89% of variation in data. Evaluating the effect of LULU curation (at 84% and 90%) for metazoans showed that LULU-curated data resolved similar ecological patterns than non-curated data, accounting from 0.5% (COI) to 1.3% (18S) of variation in data (Fig. 2).

**Figure 2.**
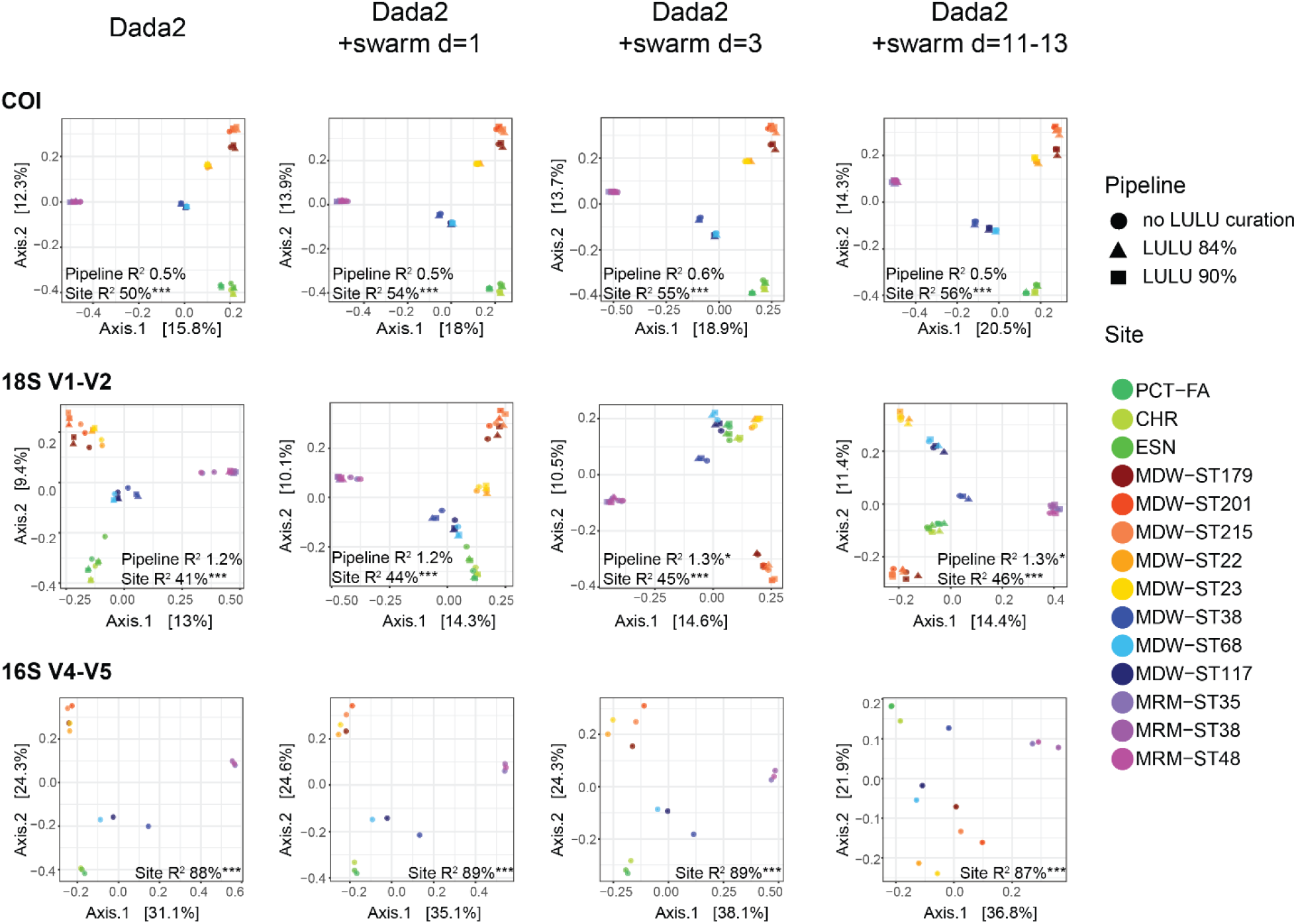
Beta-diversity patterns in ASV and OTU-centred datasets. PCoA ordinations showing community differentiation observed between sites and LULU *vs* not LULU curated samples, for the DADA2 metabarcoding pipeline with and without clustering. Metazoan datasets were clustered at *d=1-13* (COI) *d=1-11* (18S) and curated with LULU at two minimum match values. The prokaryote 16S dataset was clustered at *d=1-11*. R^2^ values and associated p-values obtained in PERMANOVAs are shown in the ordination plots. Significance codes: ***: p<0.001; **: p<0.01; *: p<0.05. Colour codes: Green: Mediterranean < 1,000 m; Red-yellow: Mediterranean-Atlantic transition zone 300-1,000 m; Blue: North Atlantic < 1,000 m; Purple: Arctic < 1,000 m.

Although ASV and OTU datasets detected similar levels of variation due to sites in PERMANOVAs, clustering levels affected the ecological patterns resolved by ordinations in rRNA loci (Fig 2). At low d values (d=1-3), ecological patterns were consistent to patterns observed in the ASV datasets, with samples segregating by site and depth. Increasing d values produced stronger segregation among sites, thus resulting in differentiation among ocean basins rather than depth. This change in resolution occurred with d values as low as d=4 for 18S but was strongest at d=11 for both rRNA loci (Fig. S3, Fig. 2).

### 4 Taxonomic assignment quality

BLAST and RDP Bayesian Classifier assignments were compared in the mock communities and natural samples, on data clustered at d=3 and curated with LULU at 84% for COI and 90% for 18S. For prokaryotes, assignment methods were compared on the ASV-level. BLAST and RDP assigned similar amounts of OTUs in the prokaryote dataset, but BLAST assigned 20-70% less OTUs in the metazoan datasets (Table S7). Assigning with BLAST at a minimum of 70% hit identity resulted in comparable results as described above. Eight of the ten species were recovered with COI and six species were recovered with 18S, while the vesicomyid bivalves were taxonomically unresolved with both loci (Fig. S4). Although most species produced one single OTU, between one and three species still resulted in 2-3 OTUs in each mock sample. Assigning the 18S dataset with RDP resulted in comparable taxonomic resolutions, although more species produced more than one OTU. Assigning the COI dataset with RDP using the MIDORI-UNIQUE database resulted in assignments of the mock samples that did not match the expected taxa and were mostly belonging to arthropods, a problem not observed with BLAST (data not shown). When the database was reduced to marine-only taxa, all 10 species were detected, and this at expected OTU abundances, once data was filtered for phylum bootstrap levels ≥ 80% (Fig S4). However, applying a phylum bootstrap minimum of 80% resulted in a strong decrease in the number of final target OTUs, particularly for COI where only 226 OTUs remained after filtering (Table S7).

This reduced recovery with RDP after applying a minimum phylum bootstrap level was not observed in prokaryotes, where 51,000-55,000 ASVs were left after filtering with both assignment methods (Table S7).

BLAST hit identities of the overall datasets varied strongly depending on phyla and marker gene (Fig. 3). For 18S, most clusters had hit identities ≥ 90%. Poorly assigned clusters (hit identity < 90%) represented less than 20% of the dataset and were mostly assigned to Nematoda, Cnidaria, Tardigrada, Porifera, and Xenacoelomorpha. For COI, nearly all clusters had similarities to sequences in databases lower than 90%. Overall, arthropods and echinoderms were detected at similar levels by both markers. The 18S barcode marker performed better in the detection of nematodes, annelids, platyhelminths, and xenacoelomorphs while COI mostly detected cnidarians, molluscs, and poriferans (Fig. 3), highlighting the complementarity of these two loci. BLAST hit identity was much higher for prokaryotes, with most clusters assigned with more than 90% similarity to sequences in databases. When datasets were filtered for RDP phylum bootstrap levels ≥ 80%, most assignments also had high genus bootstrap values for ribosomal loci. However, for COI, a considerable number of OTUs assigned to arthropods, cnidarians, molluscs, vertebrates, and poriferans still had genus bootstraps < 60%.

**Figure 3.**
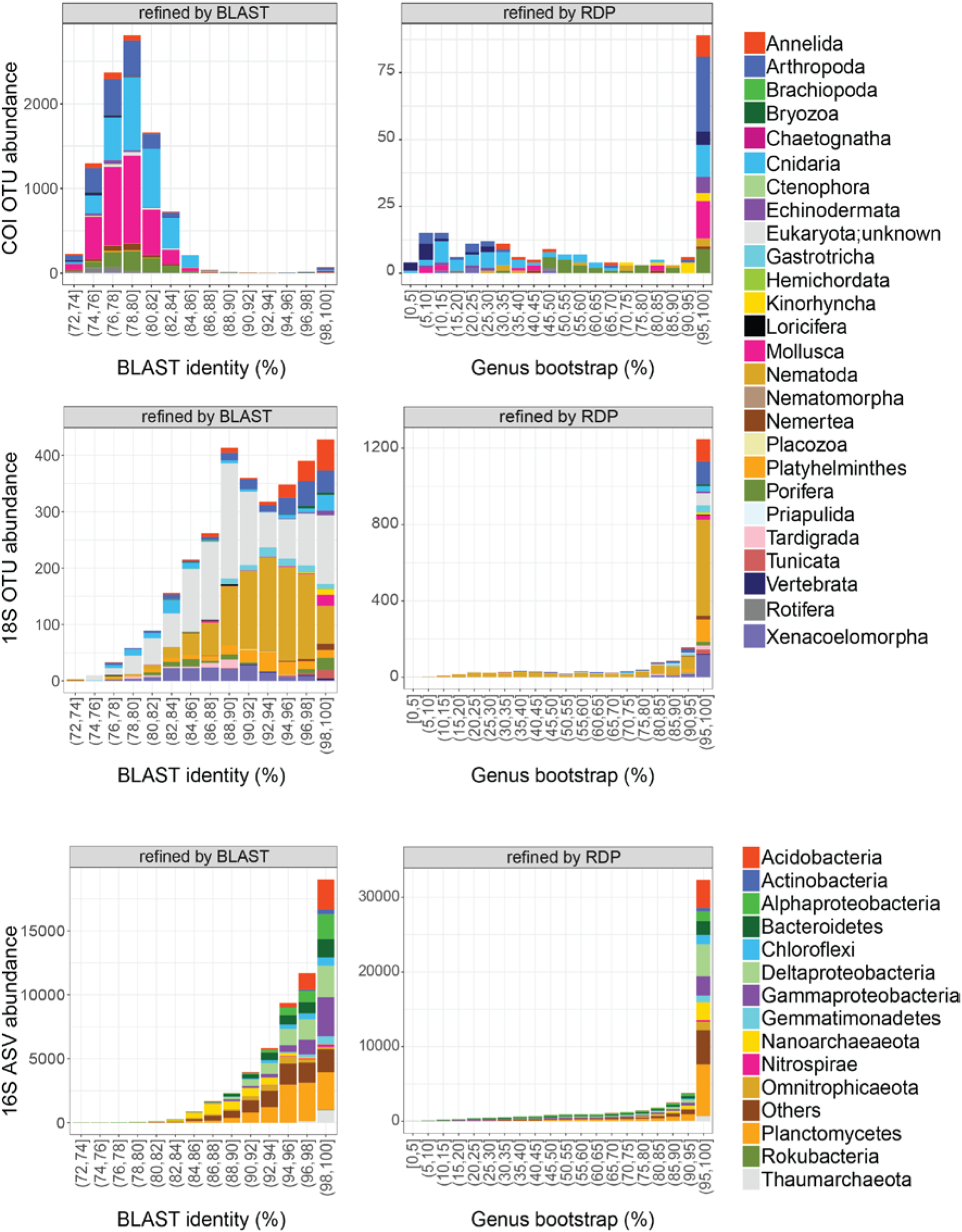
Taxonomic assignment quality of BLAST and RDP methods on metazoan and prokaryote metabarcoding datasets of 14 deep-sea sites. BLAST hit identity of all target clusters detected is given at hit identities > 70%. RDP-assigned data was filtered for phylum bootstraps ≥ 80%, and associated genus bootstraps are displayed. Taxonomic assignments were performed on the Silva132 database for 18S and 16S, and on the MIDORI-UNIQUE database, subsampled to marine taxa for COI.

## Discussion

### 1 ASVs and OTUs for genetic *vs* species diversity

The rise of HTS and the subsequent use of DNA metabarcoding have revolutionized microbiology by unlocking the access to uncultivable microorganisms, which represent by far the great majority of prokaryotes [68]. The development and improvement of molecular and bioinformatic methods to perform inventories were historically primarily developed for 16S rRNA barcode loci, before being transferred to the eukaryotic kingdom based on the use of barcode markers such as 28S and 18S rRNA, ITS, or mitochondrial markers such as COI [1], [69]. Thus, most bioinformatics pipelines were initially developed accounting for intrinsic properties of prokaryotes and concepts inherent to microbiology [70]–[72], before being transferred to eukaryotes in general, or metazoans in particular. Such application transfers require adaptations to account for differences in both concepts and basic biological features. One example is the question of the relevance of using ASVs, advocated to replace OTUs “*as the standard unit of marker-gene analysis and reporting*” [43]: an advice for microbiologists that may not apply when working on metazoans.

First, metazoans are well known to exhibit variable and sometimes very high intraspecific polymorphism in 18S-V1 and above all in COI. Second, the results on the mock samples showed that single individuals produced very different numbers of ASVs, indicating that ASV-centred datasets do not reflect actual species composition in metazoans. As this “demultiplication” will be highly variable across taxa (as seen in Fig. S2, and references such as [73] and [74]), the taxonomic compositions of samples based on ASVs will reflect genetic rather than species diversity.

Clustering ASVs into OTUs and/or curating with LULU alleviated the numerical inflation, but some species still produced more than one OTU, even at *d*-values as high as *d=11-13*. While clustering and LULU curation improved numerical results in the mock communities, they were associated with a decrease in taxonomic resolution, especially for 18S where some closely related species were merged with increasing clustering/filtering thresholds (i.e. the vesicomyid bivalves, the gastropod, and the shrimp species; Table 1). When studying natural habitats, very likely to harbour closely related co-occurring species, both LULU curation and clustering are thus likely to lead to the loss of true species diversity, particularly for low-resolution markers such as 18S. Optimal results in the mock samples, i.e. delivering the best balance between the limitation of spurious clusters and the loss of true species diversity, were obtained with LULU curation at 90% for 18S and 84% for COI, highlighting the importance of adjusting bioinformatic correction tools to each barcode marker, a step for which mock communities are most adequate.

### 2 ASVs *vs* OTUs in natural communities: adapting pipeline parameters to marker properties

Life histories of organisms, together with intrinsic properties of marker genes, determine the level of intragenomic and intraspecific diversity. Intraspecific variation is a recognised problem in metabarcoding, known to generate spurious clusters [37], especially in the COI barcode marker. Indeed, this gene region has increased intragenomic variation due to its high evolutionary rate but also due to heteroplasmy and the abundance of pseudogenes, such as NUMTs, playing an important part of the supernumerary OTU richness in COI-metabarcoding [75], [76]. Together with clustering, LULU curation at 84% proved effective in limiting the number of multiple clusters produced by single individuals, confirming its efficiency to correct for intragenomic diversity (Table 1).

The mock communities we used here did not contain several haplotypes of the same species (intraspecific variation), as is most often the case in environmental samples. This prevents us from generalizing the comparable results obtained after LULU curation of ASVs and OTUs, and the apparently limited effect of clustering in the mock samples to communities that are more complex. However, LULU curation of ASVs is not suited to account for natural haplotype diversity: not all haplotypes co-occur and when they do so, they may vary in proportion and dominance relationships, making clustering more suited to account for natural haplotypic diversity. Thus, clustering ASVs will still be necessary to produce inventories of metazoan communities that reflect species rather than gene diversity.

As expected, evaluation of clustering and LULU curation on natural samples showed distinct results for 18S and COI. Indeed, concerted evolution, a common feature of SSU rRNA markers such as 16S [68], [77] and 18S [78], limits the amount of intragenomic polymorphism. In metazoans, a lower level of diversity is expected for the slower evolving 18S gene [78], than for COI which exhibits faster evolutionary rates [79], [80]. This is reflected in the lower ASV (DADA2) to OTU (DADA2+swarm) ratios observed here for 18S (1.0-2.2.) compared to COI (2.0-2.7) data at clustering *d*-values comprised between one and seven (Table S6), underlining the different influence ‒and importance– of clustering on these loci, and the need for a versatile, marker by marker choice for clustering and curation parameters. When applying LULU to ASVs (DADA2) *versus* OTUs (DADA2+swarm) on 18S, similar cluster numbers were obtained (Fig. 1), suggesting a limited added effect of clustering for this marker once DADA2 and LULU are applied. This is in line with its slow evolutionary rate [78] leading to a limited number of haplotypes per species compared to COI. In contrast, for COI, LULU curation of the ASV dataset led to nearly twice the number of clusters (574 ± 38 at 84% and 742 ± 53 at 90%) compared to LULU curation of OTUs (at *d=6*: 334 ± 21 for 84% and 433 ± 31 for 90%). This confirms the higher intraspecific diversity of COI, and the need to combine clustering with LULU curation to account for intraspecific diversity in natural samples, especially with highly polymorphic markers such as COI.

Finally, the reproductive mode and pace of selection in microbial populations may lead to locally lower levels of intraspecific variation than the one expected for metazoans. Prokaryotic alpha diversity was however also affected by the clustering of ASVs (Fig. 1), supporting the estimation of a 2.5-fold greater number of 16S rRNA variants than the actual number of bacterial “species” [63]. The significant decrease in the number of OTUs after clustering at *d=1* (Table 2, Fig. 1, decrease of ~25%) suggests the occurrence of very closely related 16S rRNA sequences, possibly belonging to the same ecotype/species. Such entities may still be important to delineate in studies aiming for example at identifying species associations (i.e. symbiotic relationships) across large distances and ecosystems, where drift or selection can lead to slightly different ASVs in space and time, with their function and association remaining stable.

### 3 Influence on beta diversity

After focusing on alpha diversity estimates, i.e. on the numerical accuracy of inventories, the analysis of community structures showed that the LULU-curated datasets resolved similar ecological patterns as datasets not curated with LULU. However, clustering affected resolution of ecological patterns in ribosomal loci when *d* values were high, and this was not the case for COI, where similar patterns were resolved in all datasets (Fig. 2). This is in accordance with other studies reporting severe impacts of bioinformatic parameters on alpha diversity while comparable patterns of beta diversity are observed in ASV and OTU datasets, at least down to a minimum level of clustering stringency [33], [38].

Clustering and LULU curation mainly led to the decrease of the number of clusters assigned to particular taxa in both loci, such as annelids, arthropods, nematodes, or platyhelminths for 18S, and chordates, cnidarians, echinoderms, or poriferans for COI (Fig. S2). The strong decrease in cluster numbers observed in these phyla suggests that the latter have greater intraspecific polymorphism, although the decrease could also be due to the merging of closely related species, as both markers have lower taxonomic resolution in particular taxa. This has been acknowledged for 18S in general, but in nematodes in particular [81], and reported in cnidarians with COI [82].

Overall, based on alpha and beta diversity results observed in mock communities and natural samples, applying LULU at 84% seems to efficiently curate metazoan COI datasets without significant loss of species, but clustering is required, at least at *d=1*, in order to address high intraspecific polymorphism. For 18S, LULU curation seems to require values above 84% (e.g. 90%) in order to avoid the loss of species, as seen in the mock communities. However, the low taxonomic resolution obtained with this marker suggests that clustering should be performed at low *d-values* (*d<4*) to address intraspecific polymorphism without affecting beta-diversity patterns. For prokaryotes, clustering 16S ASVs at *d=1* reduces the number of detected clusters by ~25%, which may help addressing intragenomic variation when needed.

### 4 Taxonomic resolution and assignment quality

The COI locus allowed the detection of all ten species present in the mock samples, compared to seven in the 18S dataset (Table 1). This locus also provided much more accurate assignments, most of them correct at the genus (and species) level, confirming that COI uncovers more metazoan species and offers a better taxonomic resolution than 18S [83]–[85]. Our results also support approaches combining nuclear and mitochondrial markers to achieve more comprehensive biodiversity inventories [86]–[88]. Indeed, strong differences exist in amplification success among taxa [89], [90], exemplified by nematodes, which are well detected with 18S but not with COI [44]. The high complementarity of COI and 18S in terms of targeted taxa (highlighted in Fig. S2), also supports the approach taken by Stefanni et al. [91], as subsampling each gene dataset for its “best targeted phyla” and subsequently combining both seems to be a very efficient way to produce comprehensive and non-redundant biodiversity inventories.

Finally, compared to BLAST assignments, similar taxonomic resolution was observed using the RDP Bayesian Classifier on the mock samples for 18S (Fig. S4) and for COI when using the MIDORI-UNIQUE marine-only database. Poor performance of RDP using the full MIDORI database is likely due to the size of the database, and to its low coverage of deep-sea species. The problem of underrepresentation of deep-sea taxa is especially apparent with the BLAST assignments, which generally displayed low identities to sequences in databases, especially for COI (Fig. 3). Using minimum similarities of 80% for COI and 86% for 18S as cut-off values for metazoans has been shown to improve the taxonomic quality of metazoan metabarcoding datasets [91]. However, phylogenies of marine invertebrates have found high levels of species divergence (up to ~30%), even within genera [54]. Consequently, studies on deep-sea taxa have found that some invertebrate species had COI sequences diverging more than 20% from any other species present in molecular databases [55], [92]. At present, it thus seems difficult to work at taxonomic levels beyond phylum-level with deep-sea metabarcoding data when using large public databases. Small databases, taxonomically similar to the targeted communities, and with sequences of the same length as the amplified fragment of interest, are known to maximise accurate identification [93]. When using the reduced marine-only COI database, RDP provided the most accurate assignments in the mock samples when the phylum bootstrap level was ≥ 80 (Fig. S 4), although this filtering threshold drastically reduced the number of OTUs in the overall dataset (Table S7). The development of custom-built marine RDP training sets, without overrepresentation of terrestrial species, is therefore needed for this Bayesian assignment method to be effective on deep-sea datasets. With reduced and more specific databases, removing clusters with phylum bootstrap-level < 80 should be an efficient way to increase taxonomic quality of deep-sea metabarcoding datasets. At present, if accurate taxonomic assignments are sought while using universal primers, we advocate assigning taxonomy in two steps: first, using BLAST and a large database including all phyla amplifiable by the primer set, extracting the clusters belonging to the groups of interest, then re-assigning taxonomy to these target taxa using RDP and a smaller, taxon-specific database.

## Conclusions and perspectives

Using mock communities and natural samples, we evaluate several recent algorithms and assess their capacity to improve the quality of molecular biodiversity inventories of metazoans and prokaryotes. Our results support the fact that ASV data should be produced and communicated for reusability and reproducibility following the recommendations of Callahan et al. [43]. This is especially useful in large projects spanning wide geographic zones and time scales, as different ASV datasets can be easily merged *a posteriori,* and clustered if necessary afterwards. Nevertheless, clustering ASVs into OTUs will be required to obtain accurate species-level inventories, at least for metazoan communities, with a more severe influence of clustering observed on alpha diversity estimates than beta-diversity patterns. Considering 16S polymorphism observed in prokaryotic species [63] and the possible geographic segregation of their populations, clustering may also be required in prokaryotic datasets, for example in studies screening for species associations (i.e. symbiotic relationships) as symbionts may be prone to differential fixation through enhanced drift [94].

Our results also demonstrated that LULU effectively curates metazoan biodiversity inventories obtained through metabarcoding. They also underline the need to adapt parameters for curation (e.g. LULU curation at 90% for 18S and 84% for COI) and clustering to each gene used and taxonomic compartment targeted, in order to identify an optimal balance between the correction for spurious clusters and the merging of closely related species.

Finally, our findings also showed that accurate taxonomic assignments of deep-sea species can be obtained with the RDP Bayesian Classifier, but only with reduced databases containing ecosystem-specific sequences.

The pipeline is publicly available on Gitlab (https://gitlab.ifremer.fr/abyss-project/), and allows the use of sequence data obtained from libraries produced by double PCR or adaptor ligation methods, as well as having built-in options for using six commonly used metabarcoding primers.

## Data accessibility

The data for this work can be accessed in the European Nucleotide Archive (ENA) database (Illumina sequencing data are available in the European Nucleotide Archive under the following projects: PRJEB33873). The dataset, including raw sequences, databases, and ASV/OTU tables, has been deposited on https://doi.org/10.12770/0b5d250b-8418-4dda-b39c-960c4481df93. Bioinformatic scripts, config files, and R scripts are available on Gitlab (https://gitlab.ifremer.fr/abyss-project/).

## Supplementary material

Supplementary tables and figures are available online: https://doi.org/10.1101/717355

## Acknowledgements

This work is part of the “*Pourquoi Pas les Abysses?*” project, funded by Ifremer and the eDNAbyss (AP2016-228) project, funded by France Génomique (ANR-10-INBS-09) and Genoscope-CEA. We wish to thank Caroline Dussart for her contribution to the early developments of bioinformatic scripts and Patrick Durand for bioinformatic support, Caroline Menguy and Stéphane Pesant for their help to sort and post data, metadata and databases, Cathy Liautard-Haag for her help with lab management, and Babett Guenther for help in barcoding. We also wish to express our gratitude to the participants and mission chiefs of the Essnaut (Marie-Anne Cambon Bonavita), MarMine (Eva Ramirez Llodra, project 247626/O30 (NRC-BI), PEACETIME (Cécile Guieu and Christian Tamburini), CanHROV (Marie Claire Fabri) and MEDWAVES (Covadonga Orejas) cruises, and Joana Ruela Boavida, Peregrino Cambeiro (Pirri), Maria Rakka and Meri Bilan for invaluable help during sampling. The MEDWAVES cruise was organised in the framework of the ATLAS Project, supported by the European Union 2020 Research and Innovation Programme under grant agreement number: 678760. We thank Stefaniya Kamenova, Tiago Pereira, and an anonymous referee for their comments on a previous version of this manuscript. Version 3 of this preprint has been peer-reviewed and recommended by Peer Community In Ecology (https://doi.org/10.24072/pci.ecology.100041)

## Author contributions

MIB and SAH designed the study, MIB and JP carried out the laboratory and molecular work; MIB and BT performed the bioinformatic and statistical analyses. LQ assisted in the bioinformatic development and participated in the study design. MIB and SAH wrote the manuscript. All authors contributed to the final manuscript.

## References

[1] A. Valentini, F. Pompanon, and P. Taberlet, ‘DNA barcoding for ecologists’, Trends in Ecology and Evolution, vol. 24, no. 2. Elsevier Current Trends, pp. 110–117, 01–Feb–2009.

[2] P. Taberlet, E. Coissac, M. Hajibabaei, and L. H. Rieseberg, ‘Environmental DNA’, Mol. Ecol., vol. 21, no. 8, pp. 1789–1793, Apr. 2012.

[3] S. Creer et al., ‘The ecologist’s field guide to sequence-based identification of biodiversity’, Methods Ecol. Evol., vol. 7, no. 9, pp. 1008–1018, 2016.

[4] M. Stat et al., ‘Ecosystem biomonitoring with eDNA: metabarcoding across the tree of life in a tropical marine environment’, Sci. Rep., vol. 7, 2017.

[5] Y. Ji et al., ‘Reliable, verifiable and efficient monitoring of biodiversity via metabarcoding’, Ecol. Lett., vol. 16, no. 10, pp. 1245–1257, 2013.

[6] N. G. Yoccoz et al., ‘DNA from soil mirrors plant taxonomic and growth form diversity’, Mol. Ecol., vol. 21, no. 15, pp. 3647–3655, 2012.

[7] D. W. Yu et al., ‘Biodiversity soup: metabarcoding of arthropods for rapid biodiversity assessment and biomonitoring’, Methods Ecol. Evol., vol. 3, no. 4, pp. 613–623, 2012.

[8] V. Slon et al., ‘Neandertal and Denisovan DNA from Pleistocene sediments’, Science (80-)., vol. 356, no. 6338, pp. 605–608, May. 2017.

[9] J. Pansu et al., ‘Environmental DNA metabarcoding to investigate historic changes in biodiversity’, Genome, vol. 58, no. 5, p. 264, 2015.

[10] A. Valentini et al., ‘Next-generation monitoring of aquatic biodiversity using environmental DNA metabarcoding’, Mol. Ecol., vol. 25, no. 4, pp. 929–942, Feb. 2016.

[11] K. Deiner, E. A. Fronhofer, E. Mächler, J. C. Walser, and F. Altermatt, ‘Environmental DNA reveals that rivers are conveyer belts of biodiversity information’, Nat. Commun., vol. 7, no. 1, p. 12544, Dec. 2016.

[12] I. Bista et al., ‘Monitoring lake ecosystem health using metabarcoding of environmental DNA: temporal persistence and ecological relevance’, Genome, vol. 58, no. 5, p. 197, 2015.

[13] T. Dejean et al., ‘Persistence of Environmental DNA in Freshwater Ecosystems’, PLoS One, vol. 6, no. 8, 2011.

[14] N. T. Evans et al., ‘Quantification of mesocosm fish and amphibian species diversity via environmental DNA metabarcoding’, Mol. Ecol. Resour., vol. 16, no. 1, pp. 29–41, 2016.

[15] V. G. Fonseca et al., ‘Second-generation environmental sequencing unmasks marine metazoan biodiversity’, Nat. Commun., vol. 1, 2010.

[16] F. Sinniger et al., ‘Worldwide analysis of sedimentary DNA reveals major gaps in taxonomic knowledge of deep-sea benthos’, Front. Mar. Sci., vol. 3, no. June, p. 92, 2016.

[17] J. W. Pawlowski et al., ‘Eukaryotic Richness in the Abyss: Insights from Pyrotag Sequencing’, PLoS One, vol. 6, no. 4, 2011.

[18] R. R. Massana et al., ‘Marine protist diversity in European coastal waters and sediments as revealed by high-throughput sequencing’, Environ. Microbiol., vol. 17, no. 10, pp. 4035–4049, 2015.

[19] C. De Vargas et al., ‘Eukaryotic plankton diversity in the sunlit ocean’, Science (80-)., vol. 348, no. 6237, 2015.

[20] G. Salazar et al., ‘Global diversity and biogeography of deep-sea pelagic prokaryotes’, Isme J., vol. 10, no. 3, pp. 596–608, 2016.

[21] G. Boussarie et al., ‘Environmental DNA illuminates the dark diversity of sharks’, Sci. Adv., vol. 4, no. 5, p. eaap9661, May 2018.

[22] H. M. Bik et al., ‘Metagenetic community analysis of microbial eukaryotes illuminates biogeographic patterns in deep-sea and shallow water sediments’. Mol. Ecol., vol. 21, no. 5, pp. 1048–59, Mar 2012.

[23] V. G. Fonseca, ‘Pitfalls in relative abundance estimation using edna metabarcoding’, Mol. Ecol. Resour., vol. 18, no. 5, pp. 923–926, Sep 2018.

[24] I. A. Dickie et al., ‘Towards robust and repeatable sampling methods in eDNA-based studies’, Molecular Ecology Resources, vol. 18, no. 5. Wiley/Blackwell (10.1111), pp. 940–952, 01–Sep–2018.

[25] C. S. Goldberg et al., ‘Critical considerations for the application of environmental DNA methods to detect aquatic species’, Methods Ecol. Evol., vol. 7, no. 11, pp. 1299–1307, 2016.

[26] P. M. Brannock and K. M. Halanych, ‘Meiofaunal community analysis by high-throughput sequencing: Comparison of extraction, quality filtering, and clustering methods’, Mar. Genomics, vol. 23, pp. 67–75, 2015.

[27] K. Deiner, J.-C. C. Walser, E. Machler, F. Altermatt, E. Mächler, and F. Altermatt, ‘Choice of capture and extraction methods affect detection of freshwater biodiversity from environmental DNA’, Biol. Conserv., vol. 183, pp. 53–63, Mar 2015.

[28] L. Zinger et al., ‘Extracellular DNA extraction is a fast, cheap and reliable alternative for multi-taxa surveys based on soil DNA’, Soil Biol. Biochem., vol. 96, pp. 16–19, 2016.

[29] R. V. Nichols et al., ‘Minimizing polymerase biases in metabarcoding’, Mol. Ecol. Resour., vol. 18, no. 5, pp. 927–939, Sep 2018.

[30] A. Alberdi, O. Aizpurua, M. T. P. Gilbert, and K. Bohmann, ‘Scrutinizing key steps for reliable metabarcoding of environmental samples’, Methods in Ecology and Evolution, 2017.

[31] G. F. Ficetola et al., ‘Replication levels, false presences and the estimation of the presence/absence from eDNA metabarcoding data’, Mol. Ecol. Resour., vol. 15, no. 3, pp. 543–556, May. 2015.

[32] E. Coissac, T. Riaz, and N. Puillandre, ‘Bioinformatic challenges for DNA metabarcoding of plants and animals’, Mol. Ecol., vol. 21, no. 8, pp. 1834–1847, 2012.

[33] N. A. Bokulich et al., ‘Quality-filtering vastly improves diversity estimates from Illumina amplicon sequencing’, Nat. Methods, vol. 10, no. 1, pp. 57–59, Jan 2013.

[34] A. M. Eren, J. H. Vineis, H. G. Morrison, and M. L. Sogin, ‘A Filtering Method to Generate High Quality Short Reads Using Illumina Paired-End Technology’, PLoS One, vol. 8, no. 6, p. e66643, 2013.

[35] A. E. Minoche, J. C. Dohm, and H. Himmelbauer, ‘Evaluation of genomic high-throughput sequencing data generated on Illumina HiSeq and Genome Analyzer systems’, Genome Biol., vol. 12, no. 11, p. R112, Nov. 2011.

[36] E. L. Clare, F. J. J. Chain, J. E. Littlefair, and M. E. Cristescu, ‘The effects of parameter choice on defining molecular operational taxonomic units and resulting ecological analyses of metabarcoding data’, Genome, vol. 59, no. 11, pp. 981–990, Nov 2016.

[37] E. A. Brown, F. J. J. Chain, T. J. Crease, H. J. MacIsaac, and M. E. Cristescu, ‘Divergence thresholds and divergent biodiversity estimates: can metabarcoding reliably describe zooplankton communities?’, Ecol. Evol., vol. 5, no. 11, pp. 2234–2251, 2015.

[38] W. Xiong and A. Zhan, ‘Testing clustering strategies for metabarcoding-based investigation of community–environment interactions’, Mol. Ecol. Resour., vol. 18, no. 6, pp. 1326–1338, Nov 2018.

[39] R. R. Sokal and T. J. Crovello, ‘The Biological Species Concept : A Critical Evaluation’, Am. Nat., vol. 104, no. 936, pp. 127–153, 1970.

[40] B. J. Callahan, P. J. McMurdie, M. J. Rosen, A. W. Han, A. J. A. Johnson, and S. P. Holmes, ‘DADA2: High-resolution sample inference from Illumina amplicon data’, Nat. Methods, vol. 13, no. 7, pp. 581–583, 2016.

[41] J. T. Nearing, G. M. Douglas, A. M. Comeau, and M. G. I. Langille, ‘Denoising the Denoisers: an independent evaluation of microbiome sequence error-correction approaches’, PeerJ, vol. 6, p. e5364, 2018.

[42] T. G. Frøslev et al., ‘Algorithm for post-clustering curation of DNA amplicon data yields reliable biodiversity estimates’, Nat. Commun., vol. 8, no. 1, 2017.

[43] B. J. Callahan, P. J. McMurdie, and S. P. Holmes, ‘Exact sequence variants should replace operational taxonomic units in marker-gene data analysis’, ISME J., vol. 11, no. 12, pp. 2639–2643, Dec 2017.

[44] A. Bucklin, D. Steinke, and L. Blanco-Bercial, ‘DNA Barcoding of Marine Metazoa’, Ann. Rev. Mar. Sci., vol. 3, no. 1, pp. 471–508, Jan 2011.

[45] J. D. Phillips, D. J. Gillis, and R. H. Hanner, ‘Incomplete estimates of genetic diversity within species: Implications for DNA barcoding’, Ecology and Evolution, vol. 9, no. 5. John Wiley & Sons, Ltd, pp. 2996–3010, 01–Mar–2019.

[46] E. Bazin, S. Glémin, and N. Galtier, ‘Population size does not influence mitochondrial genetic diversity in animals’, Science (80-)., vol. 312, no. 5773, pp. 570–572, Apr. 2006.

[47] F. Mahe, T. Rognes, C. Quince, C. De Vargas, and M. Dunthorn, ‘Swarm v2: highly-scalable and high-resolution amplicon clustering’, PeerJ, vol. 3, 2015.

[48] E. Mayr, Systematics and the origin of species, from the viewpoint of a zoologist. New York, NY: Columbia University Press, 1942.

[49] K. de Queiroz, ‘Ernst Mayr and the modern concept of species’, Proc. Natl. Acad. Sci., vol. 102, no. Supplement 1, pp. 6600–6607, May. 2005.

[50] F. M. Cohan, ‘Bacterial species and speciation’, Syst. Biol., vol. 50, no. 4, pp. 513–524, Aug 2001.

[51] D. Gevers et al., ‘Re-evaluating prokaryotic species’, Nat. Rev. Microbiol., vol. 3, no. 9, pp. 733–739, Sep 2005.

[52] M. Leray et al., ‘A new versatile primer set targeting a short fragment of the mitochondrial COI region for metabarcoding metazoan diversity: application for characterizing coral reef fish gut contents’, Front Zool, vol. 10, p. 34, 2013.

[53] A. E. Parada, D. M. Needham, and J. A. Fuhrman, ‘Every base matters: assessing small subunit rRNA primers for marine microbiomes with mock communities, time series and global field samples’, Env. Microbiol, vol. 18, no. 5, pp. 1403–1414, 2016.

[54] J. Zanol, K. M. Halanych, T. H. Struck, and K. Fauchald, ‘Phylogeny of the bristle worm family Eunicidae (Eunicida, Annelida) and the phylogenetic utility of noncongruent 16S, COI and 18S in combined analyses’, Mol. Phylogenet. Evol., vol. 55, no. 2, pp. 660–676, May. 2010.

[55] T. M. Shank, M. B. Black, K. M. Halanych, R. A. Lutz, and R. C. Vrijenhoek, ‘Miocene Radiation of Deep-Sea Hydrothermal Vent Shrimp (Caridea: Bresiliidae): Evidence from Mitochondrial Cytochrome Oxidase Subunit I’, Mol. Phylogenet. Evol., vol. 13, no. 2, pp. 244–254, Nov 1999.

[56] C. Quast et al., ‘The SILVA ribosomal RNA gene database project: improved data processing and web-based tools’, Nucleic Acids Res., vol. 41, no. D1, pp. D590–D596, Nov. 2012.

[57] R. J. Machida, M. Leray, S. L. Ho, and N. Knowlton, ‘Data Descriptor: Metazoan mitochondrial gene sequence reference datasets for taxonomic assignment of environmental samples’, Sci. Data, vol. 4, 2017.

[58] F. Escudié et al., ‘FROGS: Find, Rapidly, OTUs with Galaxy Solution’, Bioinformatics, vol. 34, no. 8, pp. 1287–1294, Apr 2018.

[59] R Core Team, ‘R: A language and environment for statistical computing.’ R Foundation for Statistical Computing, Vienna, Austria, 2018.

[60] N. M. Davis, D. M. Proctor, S. P. Holmes, D. A. Relman, and B. J. Callahan, ‘Simple statistical identification and removal of contaminant sequences in markergene and metagenomics data’, Microbiome, vol. 6, no. 1, p. 226, Dec. 2018.

[61] I. B. Schnell, K. Bohmann, and M. T. P. Gilbert, ‘Tag jumps illuminated - reducing sequence-to-sample misidentifications in metabarcoding studies’, Mol. Ecol. Resour., vol. 15, no. 6, pp. 1289–1303, Nov 2015.

[62] O. S. Wangensteen and X. Turon, ‘Metabarcoding Techniques for Assessing Biodiversity of Marine Animal Forests’, in Marine Animal Forests, S. Rossi, L. Bramanti, A. Gori, and C. Orejas Saco del Valle, Eds. Cham: Springer International Publishing, 2016, pp. 1–29.

[63] S. G. Acinas, L. A. Marcelino, V. Klepac-Ceraj, and M. F. Polz, ‘Divergence and Redundancy of 16S rRNA Sequences in Genomes with Multiple rrn Operons’, J. Bacteriol., vol. 186, no. 9, pp. 2629–2635, May. 2004.

[64] A. Y. Pei et al., ‘Diversity of 16S rRNA genes within individual prokaryotic genomes’, Appl. Environ. Microbiol., vol. 76, no. 12, pp. 3886–3897, Jun 2010.

[65] P. J. McMurdie and S. Holmes, ‘Phyloseq: An R Package for Reproducible Interactive Analysis and Graphics of Microbiome Census Data’, PLoS One, vol. 8, no. 4, p. e61217, Apr. 2013.

[66] J. Oksanen et al., ‘vegan: Community Ecology Package’. 2018.

[67] A. Baselga and C. D. L. Orme, ‘betapart : an R package for the study of beta diversity’, Methods Ecol. Evol., vol. 3, no. 5, pp. 808–812, Oct 2012.

[68] J. A. Klappenbach, P. R. Saxman, C. J. R., and T. M. Schmidt, ‘rrndb: the Ribosomal RNA Operon Copy Number Database’, Nucleic Acids Res., vol. 29, no. 1, pp. 181–184, Jan 2001.

[69] E. Bellemain, T. Carlsen, C. Brochmann, E. Coissac, P. Taberlet, and H. Kauserud, ‘ITS as an environmental DNA barcode for fungi: An in silico approach reveals potential PCR biases’, BMC Microbiol., vol. 10, p. 189, Jul. 2010.

[10] J. G. Caporaso et al., ‘QIIME allows analysis of high-throughput community sequencing data’, Nat. Methods, vol. 7, no. 5, pp. 335–336, May. 2010.

[71] P. D. Schloss et al., ‘Introducing mothur: Open-source, platform-independent, community-supported software for describing and comparing microbial communities’, Appl. Environ. Microbiol., vol. 75, no. 23, pp. 7537–7541, 2009.

[72] F. Boyer, C. Mercier, A. Bonin, Y. Le Bras, P. Taberlet, and E. Coissac, ‘OBITOOLS: a UNIX-inspired software package for DNA metabarcoding’, Mol. Ecol. Resour., vol. 16, no. 1, pp. 176–182, 2016.

[73] S. Plouviez et al., ‘Comparative phylogeography among hydrothermal vent species along the East Pacific Rise reveals vicariant processes and population expansion in the South’, Mol. Ecol., vol. 18, no. 18, pp. 3903–3917, Sep 2009.

[74] S. Teixeira et al., ‘High connectivity across the fragmented chemosynthetic ecosystems of the deep Atlantic Equatorial Belt: Efficient dispersal mechanisms or questionable endemism?’, Mol. Ecol., vol. 22, no. 18, pp. 4663–4680, 2013.

[75] D. Bensasson, D. X. Zhang, D. L. Hartl, and G. M. Hewitt, ‘Mitochondrial pseudogenes: Evolution’s misplaced witnesses’, Trends in Ecology and Evolution, vol. 16, no. 6. pp. 314–321, 01–Jun–2001.

[76] H. Song, J. E. Buhay, M. F. Whiting, and K. A. Crandall, ‘Many species in one: DNA barcoding overestimates the number of species when nuclear mitochondrial pseudogenes are coamplified’, Proc. Natl. Acad. Sci. U. S. A., vol. 105, no. 36, pp. 13486–13491, Sep 2008.

[77] J. G. Hashimoto, B. S. Stevenson, and T. M. Schmidt, ‘Rates and consequences of recombination between rRNA operons’, J. Bacteriol., vol. 185, no. 3, pp. 966–972, Feb 2003.

[78] S. Carranza, G. Giribet, C. Ribera, J. Baguñà, and M. Riutort, ‘Evidence that two types of 18S rDNA coexist in the genome of Dugesia (Schmidtea) mediterranea (Platyhelminthes, Turbellaria, Tricladida)’, Mol. Biol. Evol., vol. 13, no. 6, pp. 824–832, Jul 1996.

[79] R. J. Machida and N. Knowlton, ‘PCR Primers for Metazoan Nuclear 18S and 28S Ribosomal DNA Sequences’, PLoS One, vol. 7, no. 9, p. e46180, Sep. 2012.

[80] R. J. Machida, M. Kweskin, and N. Knowlton, ‘PCR primers for metazoan mitochondrial 12S ribosomal DNA sequences’, PLoS One, vol. 7, no. 4, 2012.

[81] S. Derycke, J. Vanaverbeke, A. Rigaux, T. Backeljau, and T. Moens, ‘Exploring the use of cytochrome oxidase c subunit 1 (COI) for DNA barcoding of free-living marine nematodes’, PLoS One, vol. 5, no. 10, p. e13716, Oct. 2010.

[82] P. D. N. Hebert, S. Ratnasingham, and J. R. de Waard, ‘Barcoding animal life: cytochrome c oxidase subunit 1 divergences among closely related species’, Proc. R. Soc. London. Ser. B Biol. Sci., vol. 270, no. suppl_1, pp. S96-9, Aug. 2003.

[83] C. Q. Tang, F. Leasi, U. Obertegger, A. Kieneke, T. G. Barraclough, and D. Fontaneto, ‘The widely used small subunit 18S rDNA molecule greatly underestimates true diversity in biodiversity surveys of the meiofauna’, Proc. Natl. Acad. Sci. U. S. A., vol. 109, no. 40, pp. 16208–12, Oct 2012.

[84] L. J. Clarke, J. M. Beard, K. M. Swadling, and B. E. Deagle, ‘Effect of marker choice and thermal cycling protocol on zooplankton DNA metabarcoding studies’, Ecol. Evol., vol. 7, no. 3, pp. 873–883, 2017.

[85] C. Andújar, P. Arribas, D. W. Yu, A. P. Vogler, and B. C. Emerson, ‘Why the COI barcode should be the community DNA metabarcode for the metazoa’, Mol. Ecol., vol. 27, no. 20, pp. 3968–3975, Oct 2018.

[86] D. A. Cowart et al., ‘Metabarcoding Is Powerful yet Still Blind: A Comparative Analysis of Morphological and Molecular Surveys of Seagrass Communities’, PLoS One, vol. 10, no. 2, p. e0117562, 2015.

[87] A. J. Drummond et al., ‘Evaluating a multigene environmental DNA approach for biodiversity assessment’, Gigascience, vol. 4, 2015.

[88] A. Zhan, S. A. Bailey, D. D. Heath, and H. J. Macisaac, ‘Performance comparison of genetic markers for high-throughput sequencing-based biodiversity assessment in complex communities’, Mol. Ecol. Resour., vol. 14, no. 5, pp. 1049–1059, 2014.

[89] P. Bhadury, M. C. Austen, D. T. Bilton, P. J. D. Lambshead, A. D. Rogers, and G. R. Smerdon, ‘Molecular detection of marine nematodes from environmental samples: overcoming eukaryotic interference’, Aquat. Microb. Ecol., vol. 44, no. 1, pp. 97–103, 2006.

[90] L. Carugati, C. Corinaldesi, A. Dell’Anno, and R. Danovaro, ‘Metagenetic tools for the census of marine meiofaunal biodiversity: An overview’, Mar. Genomics, vol. 24, pp. 11–20, Dec 2015.

[91] S. Stefanni et al., ‘Multi-marker metabarcoding approach to study mesozooplankton at basin scale’, Sci. Rep., vol. 8, no. 1, p. 12085, Dec. 2018.

[92] S. Herrera, H. Watanabe, and T. M. Shank, ‘Evolutionary and biogeographical patterns of barnacles from deep-sea hydrothermal vents’, Mol. Ecol., vol. 24, no. 3, pp. 673–689, 2015.

[93] L. Macheriotou et al., ‘Metabarcoding free-living marine nematodes using curated 18S and CO1 reference sequence databases for species-level taxonomic assignments’, Ecol. Evol., vol. 9, no. 1, pp. 1–16, Feb 2019.

[94] B. J. Shapiro, J. B. Leducq, and J. Mallet, ‘What Is Speciation?’, PLoS Genet., vol. 12, no. 3, p. e1005860, Mar. 2016.

