## Supplementary material for "A flexible pipeline combining clustering and correction tools for prokaryotic and eukaryotic metabarcoding": Supp_Tables

Table S1. Taxonomic and relative composition of the deep-sea metazoan mock communities used in this study.

| **Taxonomic group** | **Species** | **Mock 3 (%)** | **Mock 5 (%)** |
|---|---|---|---|
| Polychaeta; Eunicida | *Eunice norvegica* | 40 | 80 |
| Crustacea; Malacostraca | *Chorocaris* sp. (now *Rimicaris* sp. or *M. fortunata* ) | 3 | 0,7 |
| Crustacea; Malacostraca | *Alvinocaris muricola* | 3 | 0,7 |
| Crustacea; Malacostraca | *Munidopsis* sp. | 3 | 0,7 |
| Anthozoa;Alcyonacea | *Acanella arbuscula* | 20 | 10 |
| Anthozoa; Scleractinia;Caryophylliidae | *Desmophyllum dianthus* | 3 | 0,7 |
| Bivalvia;Veneroida; Vesicomyidae | *Calyptogena pacifica* | 3 | 0,7 |
| Bivalvia;Veneroida; Vesicomyidae | *Christineconcha regab* (formerly *Calyptogena* sp.) | 3 | 0,7 |
| Bivalvia;Veneroida; Vesicomyidae | *Vesicomya gigas* | 3 | 0,7 |
| Gastropoda; Patellogastropoda | *Paralepetopsis* sp. | 20 | 5 |

Table S2. Sampling sites and their GPS locations and associated habitats

| Station | Cruise | Depth (m) | Latitude | Longitude | Habitat | Region |
|---|---|---|---|---|---|---|
| ESN | EssNaut | 2 400 | 42,9423 | 6,7423 | Abyssal plain | Gulf of Lyon, Mediterranean |
| PCT-FA | PEACETIME | 2 800 | 37,9467 | 2,9167 | Abyssal plain | Western Mediterranean |
| MDW-ST179 | MEDWAVES | 729 | 36,4808 | -2,8945 | Seamount | Western Mediterranean |
| MDW-ST201 | MEDWAVES | 381 | 36,546 | -2,8135 | Seamount | Western Mediterranean |
| MDW-ST215 | MEDWAVES | 554 | 36,5157 | -2,7942 | Seamount | Western Mediterranean |
| MDW-ST22 | MEDWAVES | 470 | 36,5598 | -6,9492 | Mud volcano | Gibraltar Strait |
| MDW-ST23 | MEDWAVES | 470 | 36,5605 | -6,9498 | Mud volcano | Gibraltar Strait |
| MDW-ST38 | MEDWAVES | 1 920 | 36,8442 | -11,3025 | Seamount | North Atlantic |
| MDW-ST68 | MEDWAVES | 1 245 | 37,2837 | -24,7873 | Seamount | North Atlantic |
| MDW-ST117 | MEDWAVES | 1 325 | 37,34 | -24,7552 | Seamount | North Atlantic |
| MRM-ST35 | MarMine | 2 683 | 73,4643 | 7,1975 | Hydrothermal vent | Arctic |
| MRM-ST38 | MarMine | 2 684 | 73,4639 | 7,1984 | Hydrothermal vent | Arctic |
| MRM-ST48 | MarMine | 2 826 | 73,4598 | 7,2184 | Hydrothermal vent | Arctic |
| CHR | CANHROV | 2 490 | 42,7167 | 6,1333 | Marine canyon | Gulf of Lyon, Mediterranean |

Table S3. ABYSS metabarcoding pipeline.

| **Process** | **Software** | **Script(s) and command(s)** |
|---|---|---|
| Raw reads ligation-preprocessing | Abyss-preprocessing | extractR1R2.pbs using cutadapt v1.18 (-e 0.17 for 18S-V1 and 0.27 for COI , -O length of primer -1) and BBMAP Repair v38.22 |
| Read quality-filtering | Dada2 v.1.10 | filterAndTrim() in dada2main.R maxEE=2, maxN=0, truncQ=11, truncLen=220 (18S, 16S) or 200 (COI) |
| Read error learning | Dada2 v.1.10 | learnErrors() in dada2main.R nbases=1e8, multithread=TRUE, randomize=TRUE |
| Read dereplicating | Dada2 v.1.10 | derepFastq() in dada2main.R |
| Read correction | Dada2 v.1.10 | dada() in dada2main.R |
| Read merging | Dada2 v.1.10 | mergePairs() in dada2main.R minOverlap=12, maxMismatch=0 |
| Make sequence table and filter by length | Dada2 v.1.10 | makeSequenceTable() in dada2main.R seqtab[,nchar(colnames(seqtab)) %in% seq(lengthMin,lengthMax)] lengthMin= 330 (18S-V1), 300 (COI), 350 (18S-V4), 87 (18S-V9), 350 (16S) lengthMax= 390 (18S-V1), 326 (COI), 410 (18S-V4), 186 (18S-V9), 390 (16S) |
| Chimera removal | Dada2 v.1.10 | removeBimeraDenovo() in dada2main.R |
| Taxonomy assignement with RDP Classifier | Dada2 v.1.10 | assignTaxonomy () in dada2outputfiles.R minBoot=50, outputBootstraps=TRUE |
| Taxonomy assignment with BLAST+ | blastn (megablast) v.2.6.0 | blast.pbs -outfmt 11 -qcov_hsp_perc 80 -perc_identity 70 -max_hsps 1, -evalue 1e-5, then merge BLAST and RDP taxonomies using concat_blast_rdp_tax.pbs |
| Clustering (optional), chimera removal, taxonomic assignement of OTUs | FROGS v.2.0.0 | clustering.py, remove_chimera.py, affiliation_OTU_identite_couverture.py |
| Blank correction | Rscript | Data_refining.Rmd using packages decontam v.1.2.1 and phyloseq v.1.26.0 |
| Removal of unassigned and non-target clusters | | |
| Deletion of defective samples (< 10,000 target reads) | | |
| Tag-switching renormalisation | Rscript | owi_renormalize.R |
| LULU curation | LULU v.0.1 | lulu() in lulu_final.R minimum_ratio_type = "min", minimum_ratio = 1, minimum_match =84 or 90, minimum_relative_cooccurence =0.90 |

Table S4. DADA2 read-track table. Number of reads obtained in samples after each processing step. Data refining was performed in R, based on BLAST assignments obtained using the Silva training set available on the DADA2 website for 18S and 16S, and on the MIDORI database for COI.

| Sample type | Number initial of samples | Raw reads | Quality-filtered reads | Merged reads | Length-filtered reads | Non chimeric reads Dada2 | % reads retained | Number of samples after refining | Number of target reads after all refining setps |
|---|---|---|---|---|---|---|---|---|---|
| **LOCUS** | | | | | | | | | |
| **18S-V1** | | | | | | | | | |
| Control Sample | 14 | 6 141 567 | 2 508 908 | 2 441 821 | 2 200 132 | 2 186 230 | 36 | 0 | - |
| Mock Sample | 2 | 2 096 631 | 1 607 219 | 1 436 773 | 1 430 823 | 1 289 608 | 62 | 2 | 1 287 871 |
| Natural Sample | 42 | 37 590 781 | 26 828 194 | 24 826 430 | 22 636 689 | 22 297 846 | 59 | 42 | 8 872 732 |
| **COI** | | | | | | | | | |
| Control Sample | 16 | 2 146 476 | 1 053 997 | 1 024 547 | 1 015 821 | 1 015 700 | 47 | 0 | - |
| Mock Sample | 2 | 1 482 785 | 1 261 045 | 1 252 908 | 1 251 994 | 1 224 795 | 83 | 2 | 689 810 |
| Natural Sample | 40 | 31 010 653 | 26 011 238 | 25 287 002 | 22 197 457 | 22 004 407 | 71 | 40 | 6 862 596 |
| **16S-V4V5** | | | | | | | | | |
| Control Sample | 10 | 3 531 226 | 2 889 163 | 2 634 536 | 2 619 479 | 2 618 729 | 74 | 0 | - |
| Natural Sample | 42 | 12 875 651 | 9 307 729 | 7 122 154 | 7 114 195 | 6 827 513 | 53 | 42 | 6 719 153 |

Table S5. Relative read abundance (%) detected per species in the mock communities using different bioinformatic pipelines.

| 18S | Expected abundance, based on DNA input | DADA2 | DADA2+ LULU 84% | DADA2+ LULU 90% | | Expected abundance, based on DNA input | DADA2+swarm d1/d3/d4/d5/d11 | DADA2+swarm d1/d3/d4/d5/d11 + LULU 90% | DADA2+swarm d1/d3/d4/d5/d11 + LULU 84% |
|---|---|---|---|---|---|---|---|---|---|
| **Mock 3** | | | | | | | | | |
| Alcyonacea;*A.arbuscula* | 20 | 55 | 52 | 52 | Alcyonacea;*A.arbuscula* | 20 | 55/55/55/55/55 | 54/55/55/55/55 | 54/55/55/55/55 |
| Caryophylliidae;*D.dianthus* | 3 | 2 | 5 | 5 | Caryophylliidae;*D.dianthus* | 3 | 2/2/2/2/2 | 3/2/2/2/2 | 3/2/2/2/2 |
| *Alvinocaris muricola* | 3 | 1 | 1 | 2 | *Alvinocaris muricola* | 3 | 1/2/2/2/2 | 2/2/2/2/2 | 1/2/2/2/2 |
| *Chorocaris* sp. | 3 | 1 | 0 | 0 | *Chorocaris* sp. | 3 | 1/0/0/0/0 | 0/0/0/0/0 | 0/0/0/0/0 |
| *Munidopsis* sp. | 3 | 8 | 9 | 9 | *Munidopsis* sp. | 3 | 8/8/8/8/8 | 8/8/8/8/8 | 9/8/8/8/8 |
| Gastropoda;*Paralepetopsis* sp. | 20 | 0,01 | 0 | 0,01 | Gastropoda;*Paralepetopsis* sp. | 20 | 0.002/0.012/0.017/0.017/0.019 | 0.01/0.01/0.01/0.01/0.02 | 0/0/0/0/0 |
| Vesicomyidae;*P. kilmeri/C. regab/V. gigas* | 9 | 19 | 19 | 19 | Bivalvia;*P. kilmeri/C. regab/V. gigas* | 9 | 19/19/19/19/19 | 19/19/19/19/19 | 19/19/19/19/19 |
| Polychaeta;*E.norvegica* | 40 | 14 | 14 | 13 | Polychaeta;*E.norvegica* | 40 | 14/14/14/14/14 | 13/14/14/14/14 | 14/14/14/14/14 |
| Others | 0 | 0,05 | 0,04 | 0,05 | Others | 0 | 0.05/0.07/0.07/0.07/0.07 | 0.05/0.04/0.04/0.04/0.03 | 0.05/0.04/0.04/0.04/0.05 |
| **Mock 5** | | | | | | | | | |
| Alcyonacea;*A.arbuscula* | 10 | 49 | 48 | 47 | Alcyonacea;*A.arbuscula* | 10 | 49/49/49/49/49 | 48/49/49/49/49 | 49/49/50/50/50 |
| Caryophylliidae;*D.dianthus* | 0,7 | 1 | 3 | 3 | Caryophylliidae;*D.dianthus* | 1 | 1/1/1/1/1 | 2/1/1/1/1 | 2/1/1/1/1 |
| *Alvinocaris muricola* | 0,7 | 1 | 1 | 1 | *Alvinocaris muricola* | 1 | 1/1/1/1/1 | 1/1/1/1/1 | 1/1/1/1/1 |
| *Chorocaris* sp. | 0,7 | 0 | 0 | 0 | *Chorocaris* sp. | 1 | 0/0/0/0/0 | 0/0/0/0/0 | 0/0/0/0/0 |
| *Munidopsis* sp. | 0,7 | 4 | 5 | 5 | *Munidopsis* sp. | 1 | 5/4/4/4/4 | 4/4/4/4/4 | 5/4/4/4/4 |
| Gastropoda;*Paralepetopsis* sp. | 5 | 0,01 | 0 | 0,01 | Gastropoda;*Paralepetopsis* sp. | 5 | 0.005/0.007/0.007/0.007/0.007 | 0.01/0.01/0.01/0.01/0.01 | 0/0/0/0/0 |
| Vesicomyidae;*P. kilmeri/C. regab/V. gigas* | 2,1 | 12 | 11 | 12 | Bivalvia;*P. kilmeri/C. regab/V. gigas* | 2 | 12/12/12/12/12 | 12/12/12/12/12 | 11/11/11/11/11 |
| Polychaeta;*E.norvegica* | 80 | 32 | 33 | 33 | Polychaeta;*E.norvegica* | 80 | 33/32/32/33/33 | 33/33/33/33/33 | 33/33/33/33/32 |
| Others | 0 | 0,02 | 0,02 | 0,02 | Others | 0 | 0.02/0.02/0.02/0.03/0.02 | 0.02/0.01/0.01/0.01/0 | 0.02/0.02/0.01/0.01/0.01 |
| **COI** | **Expected abundance, based on DNA input** | **DADA2** | **DADA2+ LULU 84%** | **DADA2+ LULU 90%** | | **Expected abundance, based on DNA input** | **DADA2+swarm d1/d5/d6/d7/d13** | **DADA2+swarm d1/d5/d6/d7/d13 +LULU 90%** | **DADA2+swarm d1/d5/d6/d7/d13 +LULU 84%** |
| **Mock 3** | | | | | | | | | |
| *Acanella arbuscula* | 20 | 11 | 11 | 11 | *Acanella arbuscula* | 20 | 11/11/12/12/12 | 11/11/11/11/11 | 11/11/11/11/11 |
| Hexacorallia;*D.dianthus* | 3 | 4 | 4 | 4 | Hexacorallia;*D.dianthus* | 3 | 4/4/4/4/4 | 4/4/4/4/4 | 4/4/4/4/4 |
| *Alvinocaris*;*A. muricola* | 3 | 5 | 5 | 5 | Alvinocaris;*A. muricola* | 3 | 5/5/5/5/4 | 4/4/4/4/4 | 4/4/4/4/4 |
| *Chorocaris* sp. | 3 | 1 | 1 | 1 | *Chorocaris* sp. | 3 | 1/1/1/1/1 | 1/1/1/1/1 | 1/1/1/1/1 |
| Galatheidae;*Munidopsis* sp. | 3 | 15 | 15 | 15 | *Munidopsis* sp. | 3 | 15/15/15/15/15 | 15/15/15/15/15 | 15/15/15/15/15 |
| Gastropoda;*Paralepetopsis* sp. | 20 | 27 | 27 | 27 | Gastropoda;*Paralepetopsi* s sp. | 20 | 27/27/27/27/27 | 27/27/27/27/27 | 27/27/27/27/27 |
| *Phreagena kilmeri* | 3 | 8 | 8 | 8 | Bivalvia;*P. kilmeri* | 6 | 8/8/8/8/8 | 8/8/8/8/8 | 8/8/8/8/8 |
| Bivalvia;*C. regab* | 3 | 0,2 | 0,2 | 0,2 | Bivalvia;*C. regab* | | | | |
| *Vesicomya gigas* | 3 | 3 | 3 | 3 | *Vesicomya gigas* | 3 | 3/3/3/3/3 | 3/3/3/3/3 | 3/3/3/3/3 |
| Polychaeta;*E.norvegica* | 40 | 26 | 26 | 26 | *Eunice norvegica* | 60 | 26/26/26/26/26 | 26/26/26/26/26 | 27/27/27/27/27 |
| Others | 0 | 0,1 | 0,1 | 0,1 | Others | 0 | 0.01/0.02/0.02/0.02/0.02 | 0.02/0.03/0.03/0.03/0.03 | 0.04/0.03/0.03/0.03/0.03 |
| **Mock 5** | | | | | | | | | |
| *Acanella arbuscula* | 10 | 12 | 12 | 12 | *Acanella arbuscula* | 10 | 12/12/12/12/12 | 12/12/12/12/12 | 12/12/12/12/12 |
| Hexacorallia;*D.dianthus* | 0,7 | 3 | 3 | 3 | Hexacorallia;*D.dianthus* | 1 | 3/3/3/3/3 | 3/3/3/3/3 | 3/3/3/3/3 |
| *Alvinocaris*;*A. muricola* | 0,7 | 3 | 3 | 3 | Alvinocaris;*A. muricola* | 1 | 3/3/3/3/3 | 3/3/3/3/3 | 3/3/3/3/3 |
| *Chorocaris* sp. | 0,7 | 1 | 1 | 1 | *Chorocaris* sp. | 1 | 1/1/1/1/1 | 1/1/1/1/1 | 1/1/1/1/1 |
| Galatheidae;*Munidopsis* sp. | 0,7 | 9 | 9 | 9 | *Munidopsis* sp. | 1 | 9/9/9/9/9 | 9/9/9/9/9 | 9/9/9/9/9 |
| Gastropoda;*Paralepetopsis* sp. | 5 | 18 | 17 | 18 | Gastropoda;*Paralepetopsi* s sp. | 5 | 18/18/18/18/18 | 18/18/18/18/18 | 18/18/18/18/18 |
| *Phreagena kilmeri* | 0,7 | 6 | 7 | 6 | Bivalvia;*P. kilmeri* | 1 | 7/7/7/7/7 | 7/7/7/7/7 | 7/7/7/7/7 |
| Bivalvia;*C. regab* | 0,7 | 0,1 | 0,1 | 0,1 | Bivalvia;*C. regab* | | | | |
| *Vesicomya gigas* | 0,7 | 2 | 2 | 2 | *Vesicomya gigas* | 1 | 2/2/2/2/2 | 2/2/2/2/2 | 2/2/2/2/2 |
| Polychaeta;*E.norvegica* | 80 | 46 | 46 | 46 | *Eunice norvegica* | 80 | 46/46/46/46/46 | 46/46/46/46/46 | 46/46/46/46/46 |
| Others | 0 | 0,1 | 0,1 | 0,1 | Others | 0 | 0.01/0.01/0.01/0.01/0.01 | 0.01/0.01/0.01/0.01/0.01 | 0.003/0.003/0.003/0.003/0.003 |

Table S6. Number of raw, refined, and LULU-curated molecular clusters obtained for each pipeline in the three datasets.

| Locus | Pipeline | Number of raw ASVs/OTUs | Number of target ASVs/OTUs after all refining steps | Number of target OTUs after LULU 90% | Number of target OTUs after LULU 84% | ASV to OTU ratio | ASV to OTU ratio (LULU 90%) | ASV to OTU ratio (LULU 84%) |
|---|---|---|---|---|---|---|---|---|
| 18S V1-V2 | Dada2 | 57 661 | 11 304 | 3 639 | 2 132 | - | - | - |
| | Dada2+d=1 | 44 948 | 9 090 | 3 577 | 1 980 | 1,2 | 1,0 | 1,1 |
| | Dada2+d=3 | 34 569 | 6 592 | 3 084 | 1 668 | 1,7 | 1,2 | 1,3 |
| | Dada2+d=4 | 31 509 | 5 877 | 2 889 | 1 535 | 1,9 | 1,3 | 1,4 |
| | Dada2+d=5 | 28 764 | 5 249 | 2 709 | 1 446 | 2,2 | 1,3 | 1,5 |
| | Dada2+d=11 | 19 504 | 3 117 | 1 969 | 1 094 | 3,6 | 1,8 | 1,9 |
| COI | Dada2 | 78 785 | 21 663 | 17 265 | 11 987 | - | - | - |
| | Dada2+d=1 | 64 669 | 10 375 | 8 659 | 5 612 | 2,1 | 2,0 | 2,1 |
| | Dada2+d=5 | 53 749 | 8 520 | 7 428 | 4 931 | 2,5 | 2,3 | 2,4 |
| | Dada2+d=6 | 52 216 | 8 250 | 7 251 | 4 849 | 2,6 | 2,4 | 2,5 |
| | Dada2+d=7 | 50 919 | 8 009 | 7 079 | 4 732 | 2,7 | 2,4 | 2,5 |
| | Dada2+d=13 | 44 684 | 6 840 | 6 322 | 4 345 | 3,2 | 2,7 | 2,8 |
| 16S V4-V5 | Dada2 | 56 577 | 55 129 | - | - | - | - | - |
| | Dada2+d=1 | 41 746 | 40 459 | - | - | 1,4 | - | - |
| | Dada2+d=3 | 29 023 | 27 928 | - | - | 2,0 | - | - |
| | Dada2+d=4 | 25 406 | 24 341 | - | - | 2,3 | - | - |
| | Dada2+d=5 | 22 841 | 21 795 | - | - | 2,5 | - | - |
| | Dada2+d=11 | 14 631 | 13 688 | - | - | 4,0 | - | - |

Table S7. Read and cluster abundance with data refining based on BLAST and RDP taxonomy. Number of reads, and ASVs/OTUs obtained in datasets when refining was performed based on BLAST or RDP assignments (blast / rdp). Metazoan datasets (COI and 18S) were clustered (swarm with d=3) and curated with LULU at 90% for 18S and 84% for COI, while ASVs were used in the prokaryote dataset. Taxonomic affiliations were obtained using the Silva v132 database for 18S and 16S, and the MIDORI-UNIQUE database subsampled for marine taxa for COI. BLAST assignments were performed with minimal hit identity of 70% and RDP assignments with minimum phylum bootstrap at 80%.

| LOCUS | Number of raw ASVs/OTUs | % raw ASVs/OTUs assigned (BLAST / RDP) | Number of target reads after all refining steps (BLAST / RDP) | Number of final target clusters (BLAST / RDP) |
|---|---|---|---|---|
| **COI** | 57 407 | 33% / 100% | 9,219,540 / 21,594,134 | 9,455 / 226 |
| **18S V1-V2** | 34 569 | 73% / 97% | 10,549,815 / 10,194,494 | 3,084 / 1,929 |
| **16S V4-V5** | 56 577 | 97% / 95% | 6,719,153 / 6,679,356 | 55,129 / 51,474 |
