## Supplementary material for "A flexible pipeline combining clustering and correction tools for prokaryotic and eukaryotic metabarcoding": Supp_Figures

### SUPPLEMENTAL FIGURES

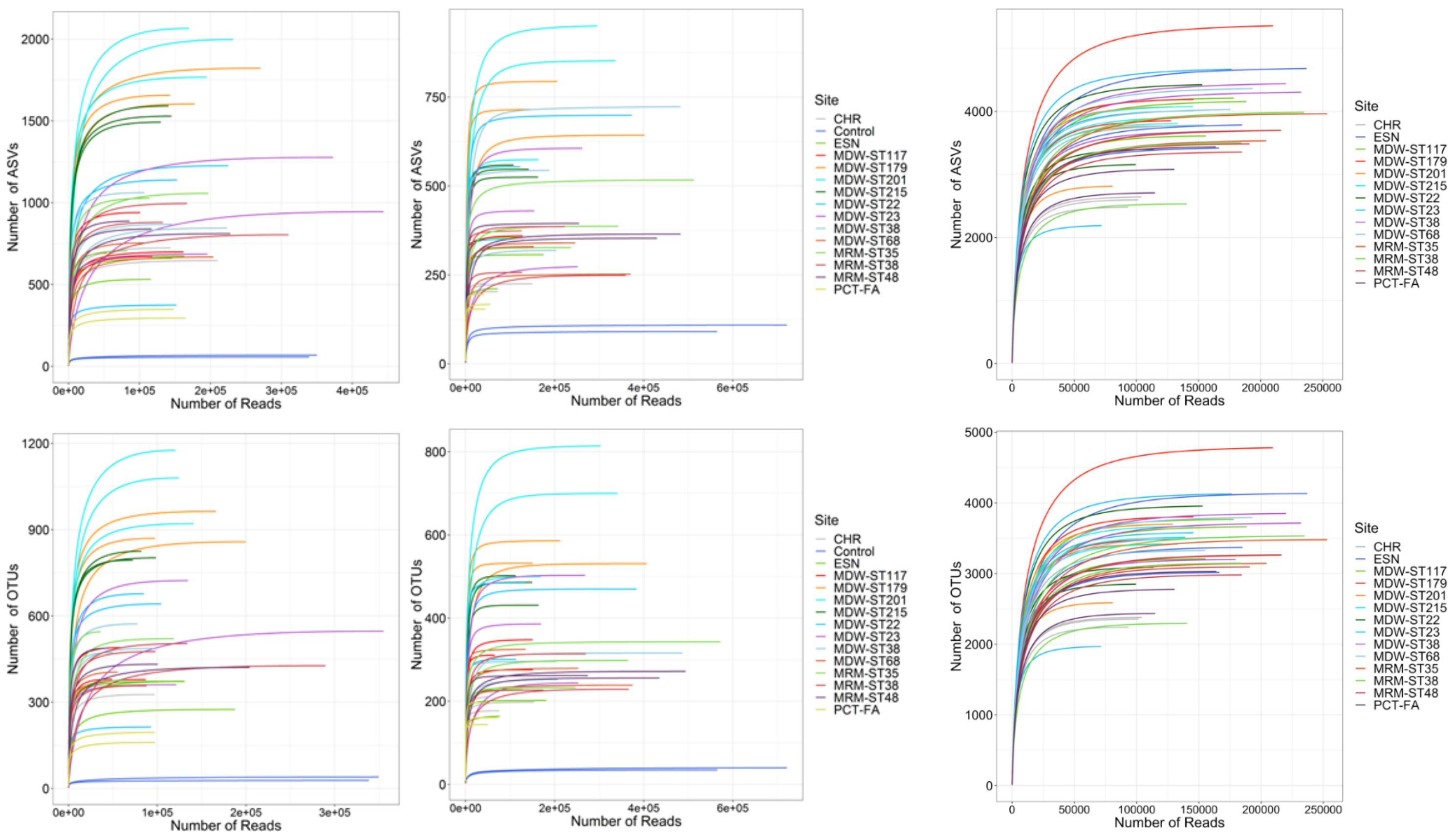


Figure S 1. Rarefaction curves in 14 deep-sea sediment sites and 2 mock control samples for metabarcoding results of the COI (left) and 18S (middle), and 16S (right) marker genes for the ASV (top) and an OTU (bottom) datasets, showing a plateau is reached in all samples.

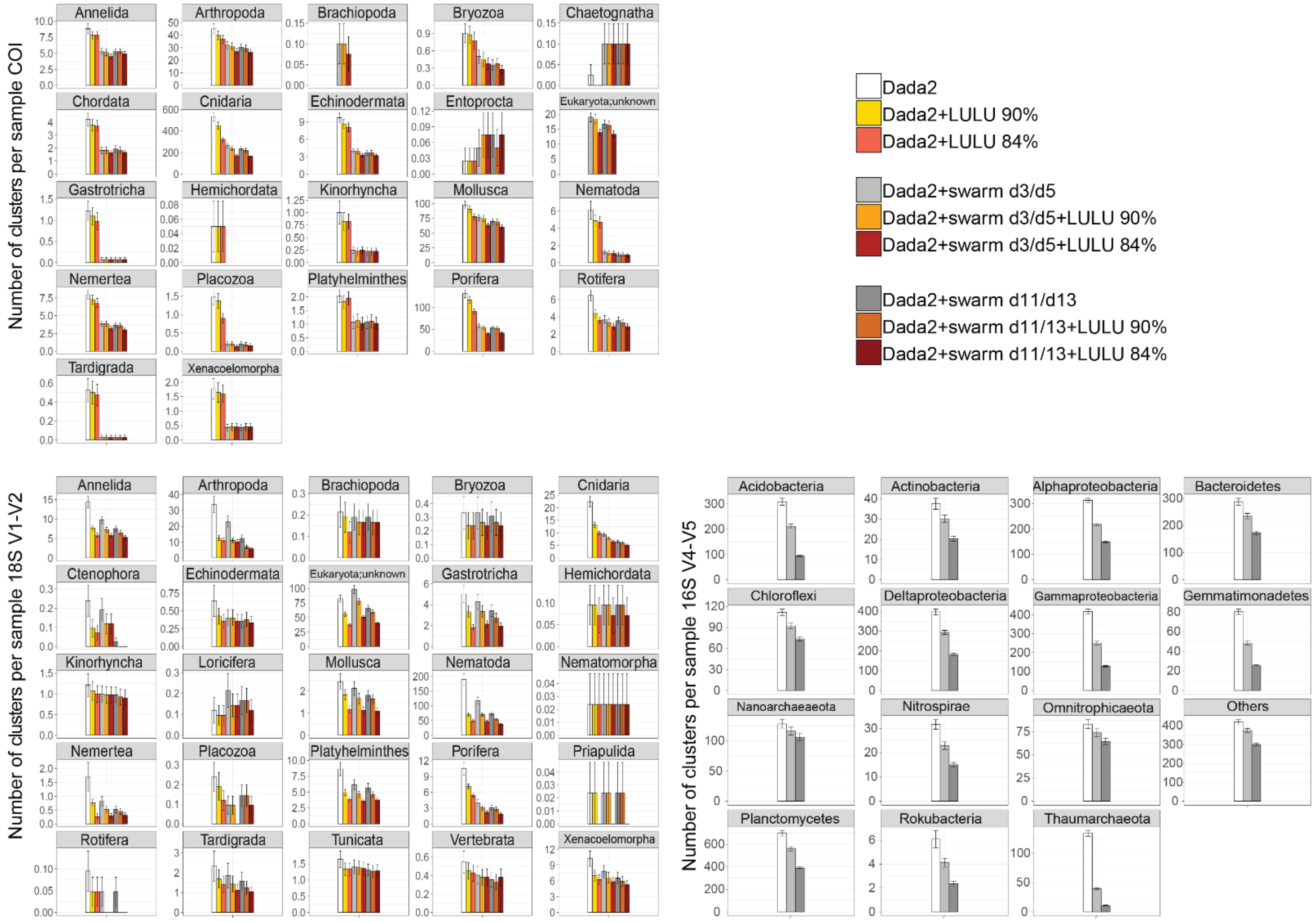


Figure S 2. Mean number of metazoan and prokaryote clusters detected per taxon in ASV vs OTU-centred datasets. Cluster numbers from sediment samples of 14 deep-sea sites were calculated from rarefied datasets. ASVs detected with the DADA2 metabarcoding pipeline are compared with OTU numbers obtained after clustering with swarm v2 (*d=3* and *d=11* for 18S and 16S, *d=5* and d=*13* for COI) and/or after curation with LULU at 84% and 90% minimum identity. Metazoans were studied using the COI and 18S marker genes, and 16S was used for prokaryotes. Error bars represent standard errors.

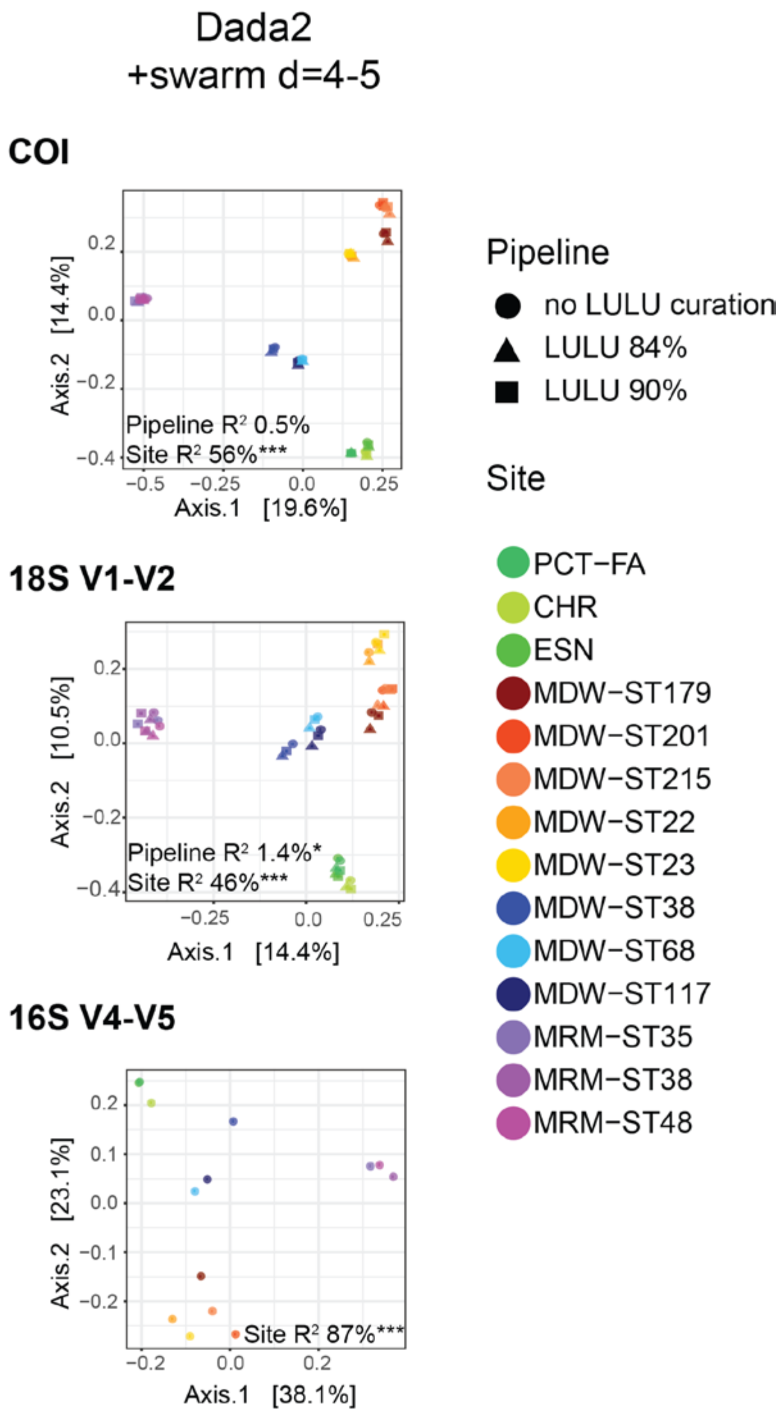


Figure S 3. Beta-diversity patterns in OTU-centred datasets at swarm clustering levels of *d=4-5*. PCoA ordinations showing community differentiation observed between sites and LULU *vs* not LULU curated samples, for the DADA2 metabarcoding pipeline with clustering at *d=4* for 18S and *d=5* for COI and 16S. $R^2$ values and associated p-values obtained in PERMANOVAs are shown in the ordination plots. Significance codes: ***:p<0.001; **:p<0.01; *:p<0.05.

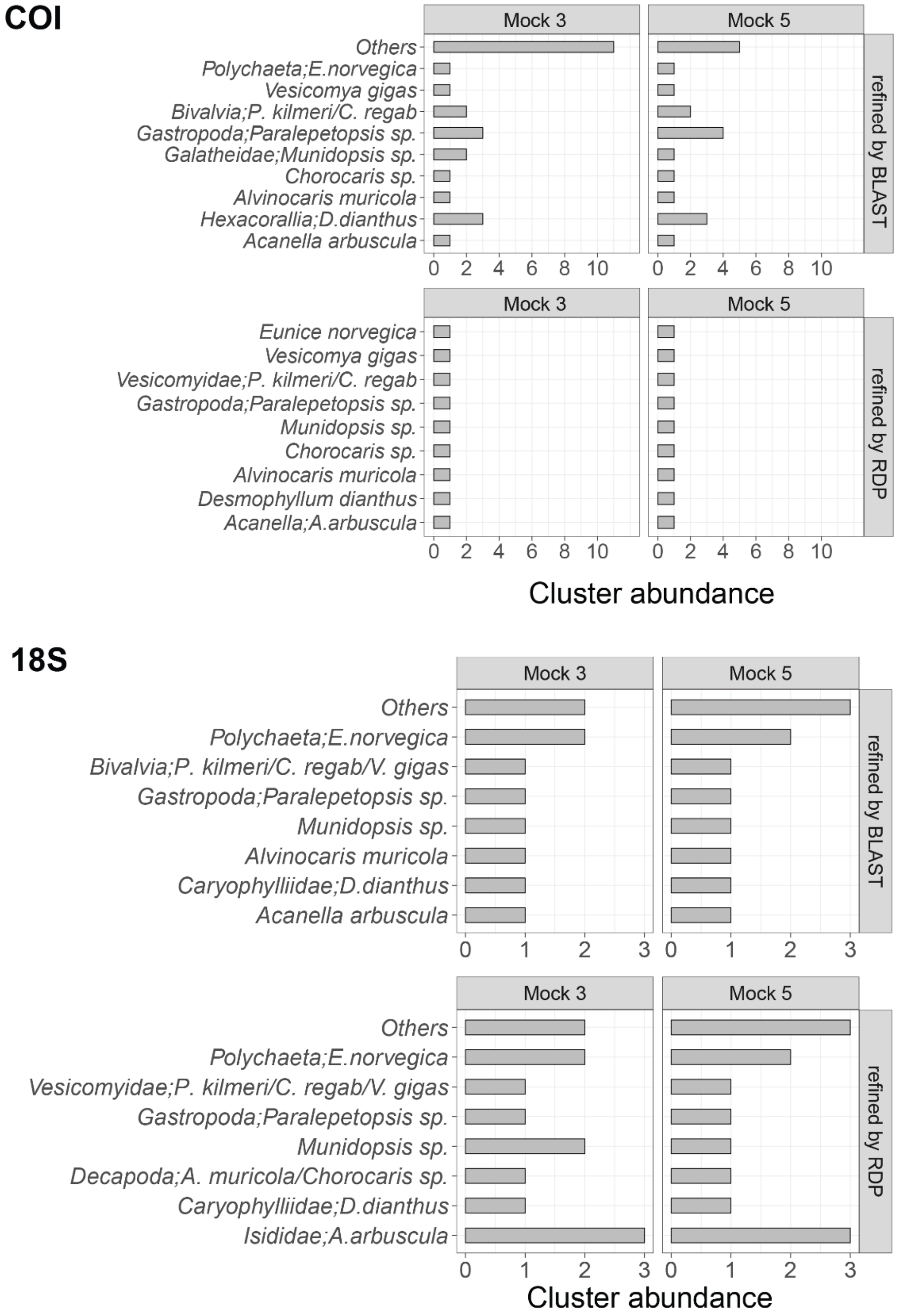


Figure S 4. Performance of RDP and BLAST taxonomic assignments methods on two mock communities of ten deep-sea species. The two mock samples were analysed within a dataset of 40-42 deep-sea samples, processed via the DADA2 metabarcoding pipeline, clustered with swarm at *d=3*, abundance-renormalized, and curated with LULU at 90% for 18S and 84% for COI. BLAST assignments were performed with a minimum hit identity of 70% (with 80% minimal coverage), and RDP assignments were kept when the phylum bootstrap was ≥ 80%. Silva132 was used as a reference database for 18S and MIDORI-UNIQUE, subsampled to marine-only taxa was used for COI. Cluster abundances were calculated on the rarefied datasets.
